# Spatial Topology Reveals Biologically Distinct Recurrent Motifs in Colorectal Cancer

**DOI:** 10.64898/2026.07.09.737584

**Authors:** Jia Yao, Yuqiu Yang, Yi Jiang, Qin Zhou, Ling Cai, Wenqi Shi, Zhikai Chi, Peiran Quan, Munir Buhaya, Bo Yao, Guanghua Xiao, Emina Huang, Yang Xie

**Affiliations:** Quantitative Biomedical Research Center, Peter O’Donnell Jr. School of Public Health, University of Texas Southwestern Medical Center, Dallas, Texas, USA; Children’s Research Institute, UTSW, Dallas, TX, USA; Department of Pathology, University of Texas Southwestern Medical Center, 5323 Harry Hines Blvd, Dallas, TX, USA; Department of Surgery, University of Texas Southwestern Medical Center, Dallas, TX, USA; Simmons Comprehensive Cancer Center, UT Southwestern Medical Center, Dallas, Texas, USA; Department of Bioinformatics, UT Southwestern Medical Center, Dallas, Texas, USA

## Abstract

Most spatial transcriptomic analyses of solid tumors focus on individual cell states or single-sample spatial domains rather than on recurrent multicellular tissue architectures shared across patients, and typically depend on predefined cell-type annotations or compartment definitions. We developed STORM (Spatial Topology analysis of Recurrent Motifs), an unsupervised graph-attention variational autoencoder that learns recurrent spatial motifs directly from cell-level graph structure and molecular profiles without cell-type labels, manual annotation, or predefined compartments. We applied to 32 Xenium sections from 16 patients with paired early-onset (EOCRC) and average-onset (AOCRC) colorectal cancer, STORM identified 10 recurrent motifs that self-organized into tumor-parenchymal, stromal, and immune macro-compartments. Among these, the Desmoplastic Fibrotic Barrier (DFB) motif, a CAF- and ECM-rich boundary architecture, was associated with restricted CD8^+^ T-cell geodesic access to tumor cores independently of CD8^+^ abundance, as demonstrated by abundance-normalized neighborhood enrichment statistics and within-sample mixed-effects models. EOCRC selectively amplified this barrier–exclusion architecture, exhibiting tighter tumor parenchyma, denser DFB shells, and a DFB-specific ECM activation program that yielded an age-specific prognostic signature in TCGA-COAD. Translation of motif macro-classes to H&E images via a Vision Transformer classifier produced an image-derived DFB-barrier composite that predicted overall survival in advanced-stage TCGA colorectal cancer. STORM provides an annotation-free framework for discovering recurrent spatial motifs and identifies a fibroblast barrier architecture whose topological association with immune exclusion is independent of effector abundance, amplified in early-onset disease, and translatable to a deployable pathology-based prognostic biomarker.

## Introduction

Colorectal cancer remains a leading cause of cancer death worldwide ^1^, and the rapid rise of early-onset disease (EOCRC, diagnosed before age 50) has strengthened the need for tissue-architectural frameworks that link tumor microenvironmental organization to clinical behavior^2,3^. The development of single-cell-resolution spatial transcriptomic platforms such as Xenium^4^, MERFISH^5,6^, VisiumHD^7^ and CosMx^8^ has made it possible to map the cellular ecosystems of solid tumors at unprecedented detail, but tools that exploit this resolution to identify recurrent multicellular tissue architectures shared across patients remain limited.

A central biological question in solid-tumor immunology is whether immune exclusion is primarily a property of cell composition or of tissue architecture. Cancer-associated fibroblast (CAF) abundance ^9^ and desmoplastic stromal reactions in CRC are known to associate with poor prognosis, but it remains unclear whether stromal geometry imposes a structural constraint on immune access to malignant cores that is independent of immune cell abundance^10–12^. Answering this question requires recurrent, cross-sample and cross-patient architectural units that explicitly preserve graph topology referred to as spatial motifs: recurrent multicellular configurations that we index at the level of each individual cell, jointly defined by that cell’s transcriptional state, the molecular profiles of its neighbors, their spatial arrangement, and reproducible across patients. So defined, a motif is an architectural label rather than a cell type, distinct also from microenvironmental regions or single-sample spatial domains.

Most current spatial analyses fall into two categories: (i) cell-state characterization, in which clusters of transcriptionally similar cells are assigned cell-type labels and then projected back onto tissue coordinates^13,14^ and (ii) spatial-domain segmentation, in which contiguous regions of similar expression are identified within individual sections.^15–17^ These approaches typically depend on predefined cell-type annotations, marker-selected compartments, or prior biological definitions of niches, and both struggle to recover multicellular configurations that recur across patients. As a result, most studies of CRC spatial biology have so far quantified the abundance and proximity of immune and stromal cells rather than the recurrent, multi-cell-type tissue architectures in which they are organized^7,18,19^.

Recent motif-oriented methods have begun to address recurrent tissue architecture more directly. TrimNN^20^ identifies cellular community motifs by modeling cell-type-labeled Delaunay triangulation graphs and detecting overrepresented topological subgraphs, whereas TissueMosaic^21^ rasterizes coarse cell-type or cell-state encodings into image patches and uses self-supervised convolutional learning to compare local tissue motifs across samples and conditions. These methods highlight the value of motif-based spatial analysis, but they remain distinct from the framework needed here: both methods presuppose either predefined cell-type labels or coarse cell-state encodings as input, and define a motif as either a discrete cell-type subgraph or a learned image-patch representation. This leaves a methodological gap between graph-based spatial-domain methods and label-dependent motif-oriented methods: an annotation-free framework that learns topology-aware, cell-level embeddings directly from gene expression and spatial graph structure and defines recurrent multicellular motifs across patients.

Here we ask whether topology-aware tissue organization adds explanatory power beyond cell abundance or composition alone. To address this, we introduce STORM (Spatial Topology analysis of Recurrent Motifs), an unsupervised graph-attention variational autoencoder. STORM learns cell-level, topology-aware embeddings directly from gene expression and spatial graph structure. It uses no cell-type labels, no manual region annotation, and no predefined tumor, stroma, or immune compartments. It defines a motif as a cluster in a shared, cross-patient latent space, so that motifs are comparable across patients. The central conceptual advance is that this topology-based view recasts immune exclusion as a property of tissue architecture rather than of cell composition. Applied to a 32-section Xenium cohort from 16 patients, STORM resolves a vocabulary of 10 motifs that self-organize, without supervision, into tumor, stromal, and immune macro-compartments. Within this map, a Desmoplastic Fibrotic Barrier (DFB) motif restricts CD8^+^ geodesic access to tumor cores independently of CD8^+^ abundance. We further show that early-onset CRC amplifies this barrier architecture, and that the framework can be read from H&E morphology alone to yield a prognostic biomarker in advanced-stage TCGA colorectal cancer.

## Results

### STORM enables topology-aware spatial motif discovery

We collected a paired Xenium spatial transcriptomics cohort comprising colorectal cancer and matched normal sections from 16 patients, including 9 EOCRC and 7 AOCRC cases (Fig. 1A, Sup Table 1). Following cell segmentation, gene-by-cell quantification, quality control, and image registration across samples, each sample retaining single-cell gene expression together with spatial coordinates.

**Figure 1.**
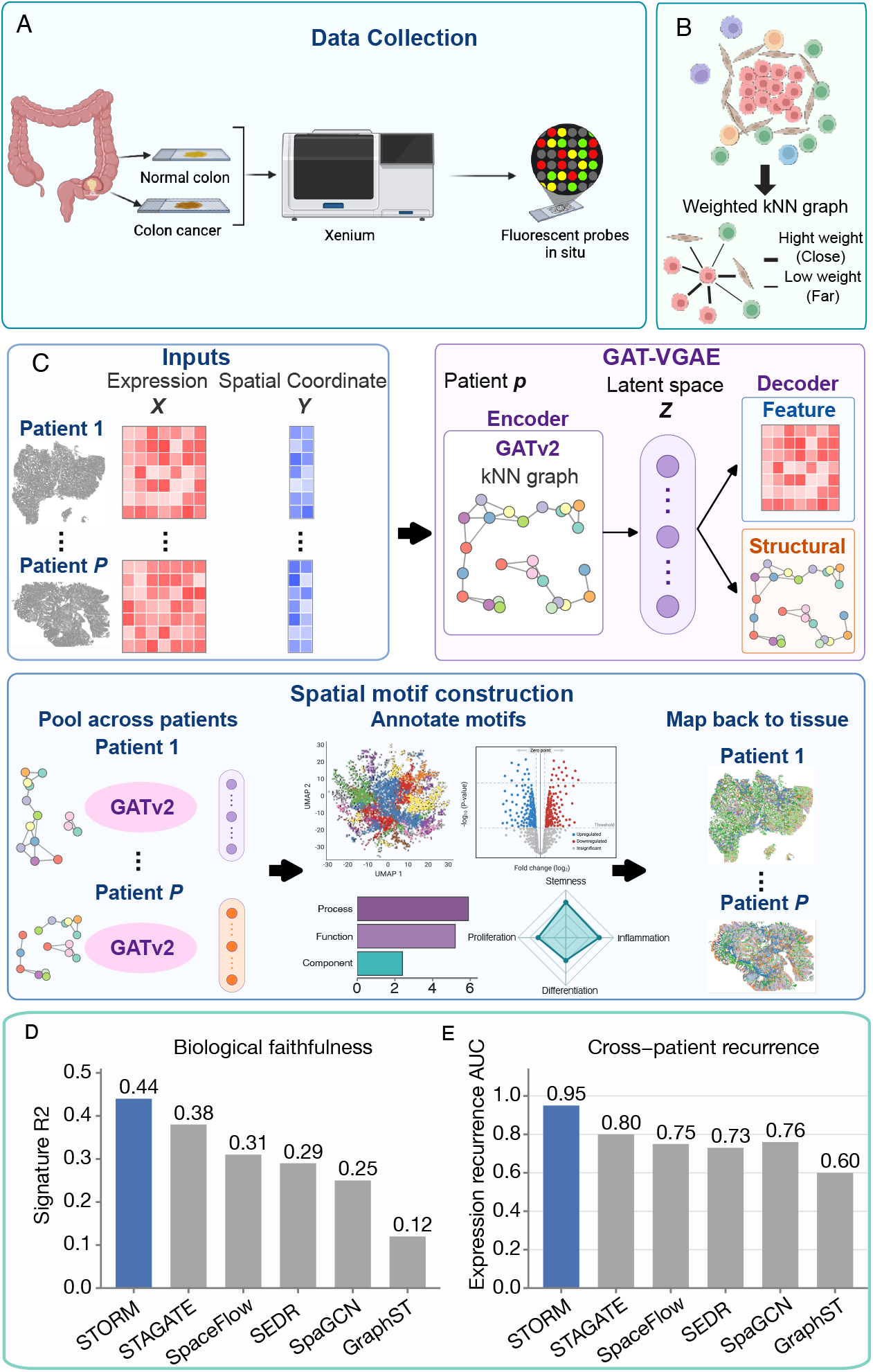
STORM, an unsupervised graph-attention variational autoencoder for spatial motif discovery. (A) Study design and data acquisition: 32 paired Xenium sections from 16 patients (9 EOCRC, 7 AOCRC), with matched tumor and normal sections per patient and matched H&E. (B) An illustration of the characterization of a spatial motif as a multicellular tissue architecture by both transcriptional states and local graph structures. (C) STORM workflow: a kNN graph is first constructed using the gene expression profile and spatial coordinates for each patient; an unsupervised graph attention variational autoencoder framework with two complementary reconstruction objectives is then trained using the kNN graphs as inputs; after training, latent embeddings from all cells are used to define spatial motifs. (D) Linear predictability of 16 predefined scanpy cell-type marker-module scores from each cell embedding. For each patient, bars summarize the mean R^2^ across modules from shuffled five-fold cross-validated ridge regression. (E) Expression-level cross-patient recurrence of globally defined K = 10 motifs. For each held-out patient, the expression recurrence AUC is the proportion of valid comparisons in which a motif’s mean standardized 375-gene expression profile is more similar to the reference profile with the same global motif label in the remaining patients than to a reference profile with a different label. This panel quantifies transcriptional-profile recurrence only. Architecture-level recurrence is shown in Figs. 2G and 3B.

Here, we define a spatial motif as a recurrent multicellular tissue architecture, indexed at the level of an individual cell and jointly characterized by the transcriptional state of that cell, the molecular profiles of its spatial neighbors, and the local graph topology connecting them (Fig. 1B). Unlike conventional cell-type labels, which assign cells by transcriptional identity alone, spatial motifs are learned from both intrinsic gene expression and local spatial context. Therefore, two transcriptionally similar cells embedded in different neighborhoods, for instance, a cancer-associated fibroblast at a desmoplastic tumor boundary versus one in a perivascular niche, receive distinct motif labels, because their graph neighborhoods differ. Conversely, cells sharing a recurrent neighborhood composition are grouped together even when their individual expression profiles diverge. Because motif labels are learned in a shared latent space across all patients, the resulting units are directly comparable across patients, tissue regions, and disease groups. We establish their recurrence at two distinct levels. At the level of transcriptional profile, each global motif label carries a consistent expression signature across patients (expression recurrence AUC; Fig. 1E). At the level of spatial architecture, the motifs obey reproducible cross-patient neighborhood-organization rules, and individual motifs show per-sample self-enrichment. Together, these results mark the motifs as reproducible organizational elements of tissue rather than sample-specific clusters or single-cell states. Applying STORM across the cohort identified multiple recurrent spatial motifs that captured shared multicellular tissue architectures across patients and could be projected back onto each section for downstream biological interpretation and validation.

To discover spatially conserved motifs, we first constructed a k-nearest-neighbour spatial graph for each section using cell coordinates, with k=10 (Fig. 1C). In this graph, each node represents a cell scaled gene expression feature, and each edge represents local spatial adjacency weighted by inverse physical distance, so that closer neighbouring cells contribute more strongly to graph message passing. To identify recurrent spatial motifs we developed an unsupervised graph attention variational autoencoder framework, termed STORM, and trained it jointly across the spatial graphs. The encoder used GATv2 attention layers to aggregate each cells’ own transcriptional state together with information from its spatial neighbors, producing a topology-aware latent embedding for every cell. A variational bottleneck regularizes the latent space, while two complementary reconstruction objectives encourage the embeddings to preserve both gene expression features and local graph structure. Specifically, the feature decoder reconstructs cell level gene expression from the embedding, whereas the structural decoder reconstructs spatial adjacency. After training, latent embeddings from all cells were clustered using mini-batch k-means to define a shared spatial motif label space. To interpret these motifs without circularity, we established each motif’s biological identity in two analytically separate steps: first from its own intrinsic transcriptional evidence including per-motif marker genes from differential expression, gene-ontology enrichment of motif-specific genes, and functional module scores for stemness, proliferation, inflammation, and differentiation. We then validated that identity against an independently derived cell-type annotation that was never provided to STORM. Because the cell-type labels were withheld from the model, their agreement with the transcriptionally defined motif identities offers a non-circular, external confirmation of biological coherence.

Since STORM is itself a graph-based representation-learning framework, we next asked whether its recurrent-motif objective could be achieved by existing graph-based spatial methods followed by the same downstream clustering procedure. We selected STAGATE^17^, Spaceflow^22^, SEDR^23^, SpaGCN^15^ and GraphST^16^, as the closest functional comparators because these methods use spatial coordinates, neighborhood graphs, or graph-representation objectives to learn spatially informed embeddings or tissue domains. Further, their embeddings can be clustered and projected back to tissue, making them appropriate baselines for testing whether general-purpose spatial graph representations are sufficient for the task addressed here. This comparison tested whether each method preserves molecular programs while producing globally defined motif labels that recur across patients. We compared STORM with these five methods using two patient-level metrics calculated from the persistent benchmark artifacts. Figure 1D measured marker-program decodability: within each patient, ridge regression was used to predict 16 Scanpy cell-type marker module scores from each embedding under shuffled five-fold cross-validation, and R^2^ values were averaged across the 16 modules. STORM had the highest mean signature R^2^ (0.442; SEM, 0.013), followed by STAGATE (0.375), SpaceFlow (0.312), SEDR (0.293), SpaGCN (0.249), and GraphST (0.124) (Fig. 1D). In paired comparisons across the patients, STORM had higher signature R^2^ than each displayed graph comparator (Holm-adjusted P ≤3.0 × 10^−4^; paired Cohen’s d_z_ = 1.26-4.49). Figure 1E measured within-cohort cross-patient consistency of the K = 10 motif expression profiles. For each held-out patient, the expression recurrence AUC quantified whether a motif’s mean standardized 375-gene expression profile was more similar to the profile with the same global motif label in the remaining patients than to profiles with other labels. This metric establishes recurrence at the level of transcriptional profile only. In isolation, the expression recurrence AUC does not test whether a motif’s spatial architecture recurs. Architecture-level recurrence is established separately. It rests on the reproducibility of the motif × motif neighborhood-enrichment structure across patients and on per-sample motif self-enrichment (Fig. 2G). STORM again had the highest mean expression recurrence AUC (0.951; SEM, 0.014), compared with 0.805 for STAGATE, 0.757 for SpaGCN, 0.745 for SpaceFlow, 0.727 for SEDR, and 0.603 for GraphST (Fig. 1E), and its patient-level AUC was higher than that of each displayed comparator (Holm-adjusted P ≤ 0.00231; d_z_ = 1.51-3.28). This performance gap is consistent with STORM’s original motivation. Recurrent motif discovery requires embeddings that retain molecular programs, encode local graph topology, and remain comparable across patients. STORM addresses these needs through feature reconstruction, structural adjacency reconstruction, distance-aware graph attention, and joint cross-patient training. The benchmark therefore suggests that STORM’s task-specific architecture is better suited to recovering recurrent, molecularly interpretable, topology-aware tissue motifs that can be compared across patients and used in downstream analyses.

**Figure 2.**
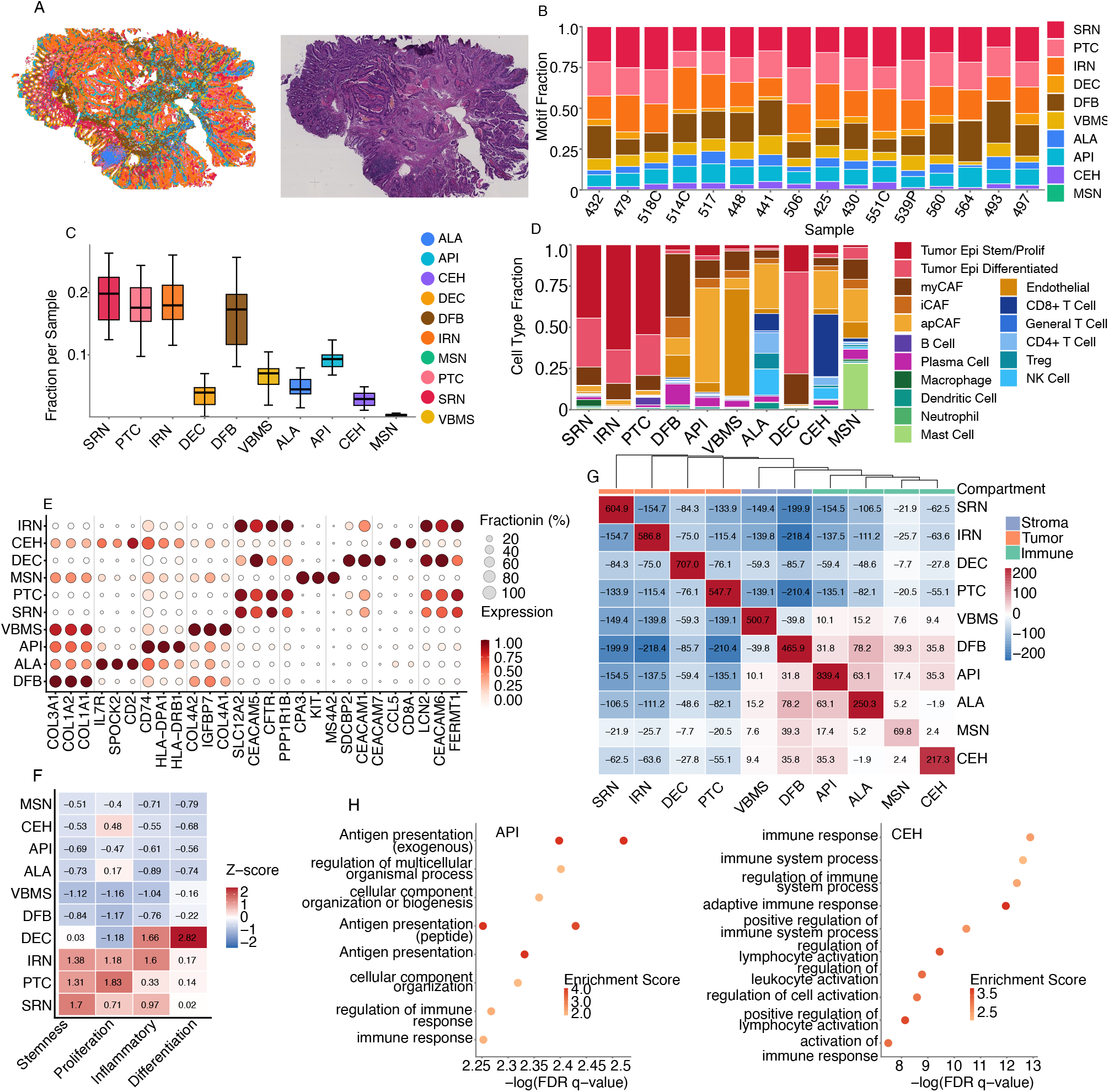
A ten-motif spatial atlas resolves biologically distinct cellular architectures across tumor, stromal, and immune compartments of colorectal cancer. (A) Representative CRC section colored by ten STORM motifs (left) with the matched H&E image (right). (B) Per-sample motif fractions across the 16 cancer sections (stacked bar, top). (C) Sample-fraction boxplots across motifs. (D) Cell-type composition per motif computed from independent cell-type annotation; each motif shows a distinct multicellular composition (e.g., DFB = myCAF/iCAF/apCAF-dominated; CEH = CD8-dominated; ALA = mixed T/B/plasma cells; API = macrophage/monocyte/dendritic-cell-dominated). (E) Top marker gene dot plot for the ten motifs, with motif-specific marker blocks along the diagonal. (F) Functional module scores (Stemness, Proliferation, Inflammatory, Differentiation) for each motif, highlighting IRN as a triple-positive injury/regeneration state and DEC as inflammatory despite differentiation. (G) Global motif × motif neighborhood-enrichment Z heatmap with hierarchical clustering recovering tumor (SRN, IRN, DEC, PTC), stromal (VBMS, DFB), and immune (API, ALA, MSN, CEH) blocks without supervision; the compartment annotation bar at the top of the dendrogram is derived from the data and not from any cell-type label. (H) Gene Ontology enrichment bubble plots for API (left, antigen presentation of exogenous antigen and MHC class II processing) and CEH (right, adaptive immune response, lymphocyte activation, and positive regulation of T-cell activation), illustrating motif-specific biological identity.

### A 10-motif atlas reveals tripartite organization of the colorectal cancer microenvironment

The above benchmarks show that STORM yields internally consistent, cross-patient motif labels and embeddings from which marker programs are more readily decoded. However, quantitative consistency does not establish what the motifs represent. Applying the two-step interpretation framework across the cohort, STORM resolved 10 recurrent spatial motifs (Sup Table 1; Figure 2A) that recurred across the 16 cancer sections (Figure 2B,C).

For every motif, the two independent lines of evidence converged: intrinsic transcriptional signature including marker genes (Figure 2E), Gene ontology enrichment (Figure 2H), and functional module scores (Figure 2F) were assigned a coherent identity that was then corroborated by its independently derived cell-type composition (Figure 2D). Because the cell-type annotation was withheld from STORM, this convergence both justifies each motif’s identity and confirms that the motifs are biologically coherent multicellular architectures rather than statistical artefacts, a conclusion further supported by stability analyses excluding batch effects and patient differences (Sup. File 1 and Sup. Fig. 1A-H). On this combined evidence, the 10 motifs organize into tumor-parenchymal, stromal, and immune compartments, which we describe below before considering their global spatial organization.

#### Tumor parenchyma

Four motifs constituted the tumor parenchyma (Sup Table 2), organized along a regeneration-proliferation-differentiation axis (Sup Table 1; Figure 2D-F). The Stem-Like Regenerative Niche (SRN; Motif 4) carried the highest stemness score with only moderate proliferation, marking a slow-cycling self-renewing reservoir, whereas the Proliferative Tumor Core (PTC; Motif 5) showed the atlas’s highest proliferation program and drove bulk expansion. Most distinctive was the Inflammatory Regenerative Niche (IRN; Motif 9), uniquely triple-positive for stemness, proliferation, and inflammation — a profile consistent with an injury-induced dedifferentiation/regeneration program rather than passive inflammation^24^. The Differentiated Epithelial Compartment (DEC; Motif 7), the only motif dominated by differentiated tumor epithelium, combined a terminal-differentiation program with prominent *IL1B* expression, identifying it as a non-canonical epithelial source of pro-inflammatory signaling^25^. Cell-type composition confirmed all four as tumor-epithelial (Figure 2D).

#### Stromal architecture

Two motifs formed the stromal architecture (Sup Table 2). The Desmoplastic Fibrotic Barrier (DFB; Motif 0) was defined by fibrillar collagens (*COL1A1/2, COL3A1*) with GO enrichment for collagen fibril and extracellular-matrix organization and cell adhesion, and comprised myofibroblastic, inflammatory, and antigen-presenting CAFs (myCAF, iCAF, apCAF), establishing its desmoplastic identity (dissected in Figure 3). The Vascular Basement Membrane Scaffolding (VBMS; Motif 3), defined by basement-membrane collagens (*COL4A1/2*), *IGFBP7*, and *RGS5* with angiogenesis enrichment and composed of endothelial and pericyte populations^26^, marked the perivascular scaffold of the tumor microenvironment (TME).

**Figure 3.**
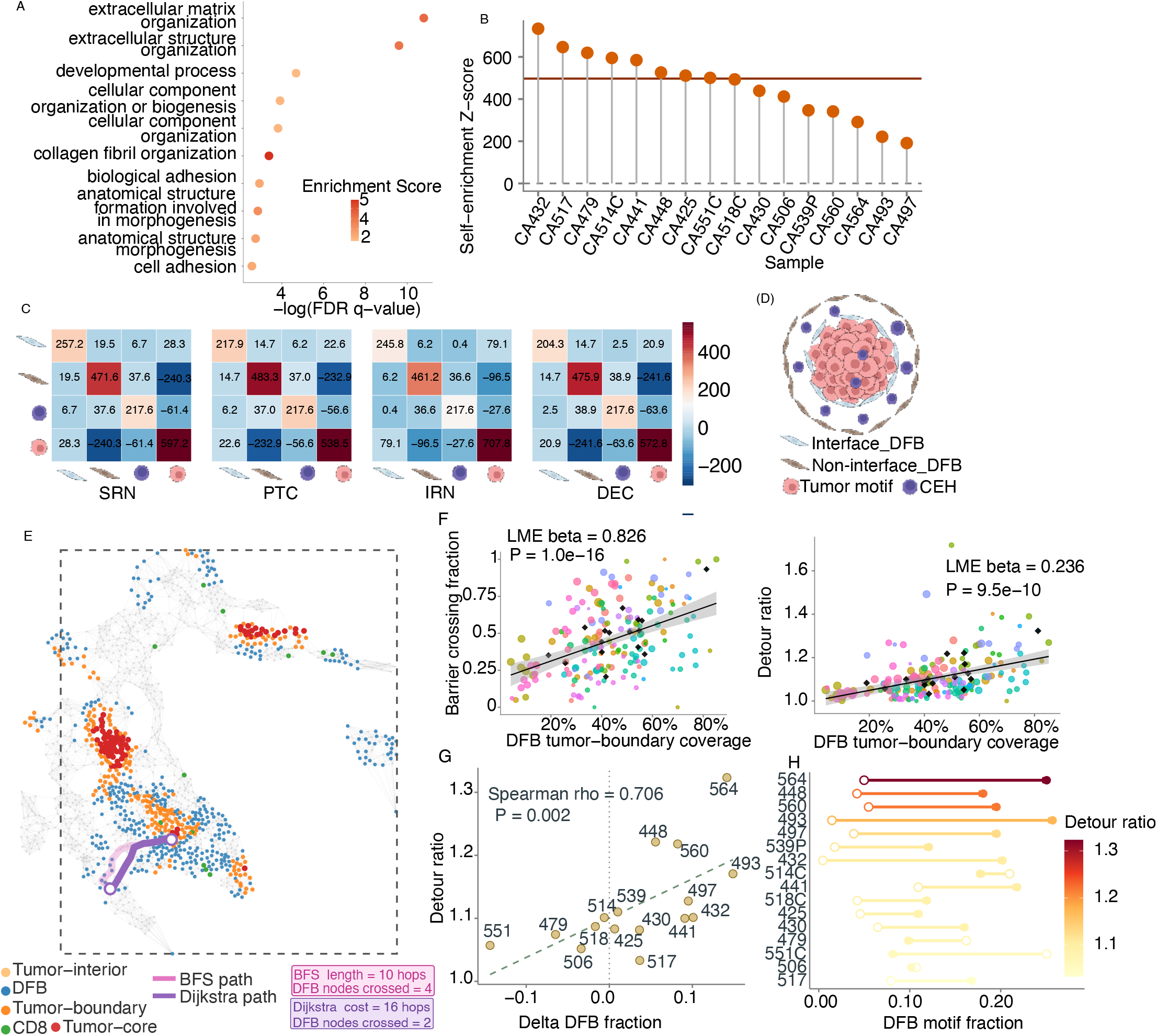
The Desmoplastic Fibrotic Barrier (DFB) motif restricts CD8 geodesic access to tumor cores independently of immune abundance. (A) GO enrichment of DFB-specific marker genes; leading terms: collagen fibril organization, ECM organization, cell adhesion. (B) Per-sample DFB self-enrichment Z scores (n = 16; range 191-734, median ≈ 500; Fisher combined one-sided P = 5.14 × 10^−23^). (C) For each tumor motif (SRN, PTC, IRN, DEC), a 4 × 4 neighborhood-enrichment Z matrix on interface DFB, non-interface DFB, CEH, and the tumor motif (aggregated, n = 16). Interface DFB is co-enriched with tumor (Z = +20.9 to +79.1); non-interface DFB and CEH are excluded from tumor; CEH co-enriches with non-interface DFB (Z ≈ +37) but not interface DFB. Z is abundance-normalized. (D) Spatial-layout schematic from (C): tumor core → thin interface-DFB band → bulk non-interface-DFB layer trapping CEH cells, forming a thin shell that separates cytotoxic effectors from malignant cells. (E) Geodesic-access pipeline: cell-level KNN graph; tumor-boundary (≥1 non-malignant neighbor) and tumor-core (≥3 hops interior) nodes; unweighted versus DFB-penalty-weighted shortest paths from each CD8 cell to its nearest core node, yielding a per-cell barrier-crossing indicator and detour ratio (weighted ÷ unweighted). (F) Tile-level linear mixed-effects model (sample as random effect; n = 182 tiles, 16 samples): DFB boundary coverage predicts barrier-crossing fraction (β = 0.826, P = 1.0 × 10^−16^), detour-ratio mean (β = 0.236, P = 9.5 × 10^−10^), and median (β = 0.166, P = 1.6 × 10^−5^). Sample-level Spearman: barrier crossing ρ = 0.503 (P = 0.047); detour ratio ρ = 0.618 (P = 0.011). Sample-level and penalty-robustness analyses in Sup Fig 2. (G) Patient-level association between tumor-induced DFB expansion (ΔDFB fraction relative to matched-normal baseline) and CD8 core detour ratio; each point is one paired specimen (Spearman ρ = 0.706, P = 0.0022, n = 16). Dotted line, no net expansion; dashed line, linear trend. (H) Within-patient DFB fraction shifts from matched normal (open circles) to cancer (filled circles); arrows show paired change. Patients are ordered by increasing CD8 detour ratio (color scale); higher detour ratios show larger positive DFB shifts.

#### Immune interface

Four motifs made up the immune interface (Sup Table 2). Central to the immune-exclusion story below, the Cytotoxic Effector Hub (CEH; Motif 8) carried by far the strongest GO enrichment for adaptive immune response and T-cell activation and was dominated by CD8^+^ T and NK cells, defining the cytotoxic killing zone. It was complemented by the Antigen-Presenting Interface (API; Motif 2; MHC-II^+^, macrophage/monocyte/DC-rich) and the Adaptive Lymphoid Aggregate (ALA; Motif 1), a mixed T/B/plasma compartment consistent with (pre-)tertiary lymphoid structures^27^. The rarest motif, the Mastocytic Sentinel Niche (MSN; Motif 6; 0.3%), was transcriptionally extreme (*CPA3, KIT, MS4A2*) and sat at the ECM-immune boundary, consistent with a tissue-resident sentinel role^28^.

#### Tripartite organization of the motif atlas

Having defined each motif individually from its transcriptional evidence, we next asked whether the motifs also obey reproducible spatial organizational rules which is information that marker identity alone cannot provide. We analyzed the mean motif × motif neighborhood-enrichment Z matrix, which encodes only spatial adjacency between motifs and is independent of the marker and module-score evidence used above to name them, and two distinct kinds of information emerged. As cross-modal corroboration, unsupervised hierarchical clustering of this purely spatial matrix partitioned the ten motifs into the same three groups suggested by their transcriptional identities — tumor parenchyma (SRN, IRN, DEC, PTC), stromal architecture (VBMS, DFB), and immune interface (API, ALA, MSN, CEH) (Figure 2G, top dendrogram and annotation bar). Because the Z matrix never accessed the marker biology, this agreement is independent evidence that the transcriptionally defined compartments are spatially coherent tissue units rather than expression categories alone.

Beyond the grouping, the off-diagonal structure revealed adjacency rules not recoverable from motif identity that directly motivate the immune-exclusion analyses below. Strong diagonal self-enrichment (combined mean Z 73-694) showed each motif aggregates into spatially coherent patches. Within the tumor compartment, the substates were mutually exclusive at the single-edge scale (SRN ↔PTC, SRN ↔DEC, SRN ↔IRN all Z < −80; SRN the most insulated “clonal island”), so tumor parenchyma forms a macro-compartment through shared external relations despite internal spatial segregation. The desmoplastic compartment was sharply segregated from tumor at the single-hop scale (DFB ↔SRN ≈−190, DFB ↔IRN ≈ −201, DFB ↔PTC ≈ −187). Most informatively, cytotoxic effectors tracked the stroma rather than the tumor: CEH ↔DFB was positive (+43.8 in EOCRC, +26.1 in AOCRC) while CEH ↔tumor edges were negative — a topological signature that cytotoxic cells accumulate at the stromal interface rather than within parenchyma, the spatial premise dissected in Figure 3. As a sanity check, the canonical antigen-presentation pair ALA ↔API was positive (+76.6 in EOCRC, +62.3 in AOCRC), recovered without supervision.

Together, these results show that the CRC microenvironment is spatially modular: an unsupervised, adjacency-based analysis of STORM motifs independently reproduces the tripartite compartmentalization and, more importantly, exposes the segregation and interface rules, tumor-substate exclusivity, DFB-tumor segregation, and CEH accumulation at the stromal interface that frame the barrier-exclusion architecture examined next.

#### The Desmoplastic Fibrotic Barrier is a recurrent boundary-layer motif associated with restricted CD8^+^ geodesic access to tumor cores

The tripartite analysis raised a specific, testable form of the question posed in the Introduction: whether immune exclusion reflects cell composition or tissue architecture. Among the ten motifs, DFB was the only stromal motif positioned at the boundary between the tumor and immune compartment: DFB segregated sharply from tumor parenchyma yet co-localized with the cytotoxic effector hub (CEH), exactly the topology expected if stromal architecture, rather than a shortage of effector cells, separated cytotoxic cells from malignant cores. We therefore focused on DFB to test whether its geometry imposes a structural constraint on CD8^+^ access that is independent of CD8^+^ abundance. Throughout this section we keep two label layers distinct: STORM motif labels (e.g., DFB, CEH), which mark cells by their neighborhood architecture and not by cell type, and the independent cell-type annotation (e.g., CD8^+^ T cells). The spatial blueprint below is built from motif labels; the geodesic quantification is grounded in the annotated CD8^+^ T cells.

DFB formed reproducible, spatially coherent units across patients. GO enrichment of DFB-specific genes recovered collagen fibril organization, extracellular matrix organization, and cell adhesion as the leading terms (Figure 3A), consistent with a CAF- and ECM-rich stromal architecture, and per-sample DFB self-enrichment Z scores ranged from 191 to 734 (median ≈ 500), with every sample significantly enriched (Fisher combined one-sided p = 5.14 × 10^−23^; Figure 3B).

STORM discovers DFB as a single motif. The next partition is not a STORM motif. Instead, we specify a downstream, investigator-defined split of that motif’s cells, based on tumor adjacency. To resolve how DFB is positioned relative to tumor and to cytotoxic effectors, we divided DFB-motif cells into two sub-populations. Interface DFB cells are DFB-motif cells in direct contact with a tumor motif. Non-interface DFB cells are the remaining DFB-motif cells. They form the bulk fibrosis at greater distance from the tumor. For each of the four tumor motifs (SRN, PTC, IRN, DEC) we computed a 4 × 4 neighborhood-enrichment Z matrix over interface DFB, non-interface DFB, CEH, and the tumor motif, aggregated across 16 samples (Figure 3C). Three patterns recurred across all four tumor contexts. First, interface DFB was positively associated with the tumor motif (Z = +20.9 in IRN to +79.1 in DEC; Fisher combined p ≤ 5.9 × 10^−17^), confirming that this band directly abuts tumor. Second, both non-interface DFB (Z = −96.5 in DEC to −241.6 in IRN) and CEH (Z = −27.6 in DEC to −63.6 in IRN) were strongly excluded from tumor, so neither the bulk fibrosis nor the cytotoxic-effector motif penetrates the parenchyma. Third, and most informatively, CEH was reproducibly co-enriched with non-interface DFB (Z = +36.6 to +38.9; Fisher combined p down to 6.5 × 10^−17^) but essentially uncorrelated with interface DFB (Z = +0.4 to +6.7). Because each Z value compares observed adjacency to a permutation null that preserves per-motif cell counts, this co-enrichment holds independently of CEH abundance — a first, abundance-free line of evidence that the cytotoxic-effector motif is held in the bulk stroma rather than at the tumor-facing interface.

These adjacencies assemble a concrete spatial blueprint (Figure 3D): moving outward from the tumor, a parenchymal core is wrapped by a thin interface-DFB band in direct tumor contact, then a bulk non-interface-DFB layer in which the CEH motif preferentially resides, then the remaining microenvironment. The cytotoxic-effector compartment is therefore not absent from the tissue; it is held within the bulk stroma, separated from malignant cells by a thin but circumferentially continuous interface-DFB band.

We then grounded this topology in the annotated immune compartment by modeling CD8^+^ T-cell trajectories on the per-section spatial graph (Figure 3E). Here the analysis moves from motif labels to the independent cell-type annotation: tumor-boundary and tumor-core nodes were defined from annotated malignant cells (boundary, ≥1 non-malignant neighbor; core, ≥3 graph steps interior), the migrating agents were annotated CD8^+^ T cells, and DFB-motif nodes served as the structural barrier. For each CD8^+^ T cell we computed (i) the unweighted shortest path to the nearest tumor-core node, recording whether it crossed ≥1 DFB node (barrier-crossing fraction), and (ii) a DFB-penalty-weighted shortest path, yielding a detour ratio (weighted ÷ unweighted). Per-cell statistics were summarized per tile and per sample; tile-level analyses used a linear mixed-effects model with sample as a random effect, and sample-level analyses used Spearman correlations. Across 182 tiles from 16 samples, DFB boundary coverage strongly predicted impaired CD8^+^ access (Figure 3F): barrier-crossing fraction β = 0.826 (p = 1.0 × 10^−16^), detour-ratio mean β = 0.236 (p = 9.5 × 10^−10^), and detour-ratio median β = 0.166 (p = 1.6 × 10^−5^). At the sample level, DFB boundary coverage correlated with CD8^+^ barrier-crossing fraction (Spearman ρ = 0.503, p = 0.047) and detour ratio (ρ = 0.618, p = 0.011), matching the mixed-effects direction and rank order, and the tile-level effect was stable across DFB-traversal penalties (1, 3, 5, 8; Sup Fig 2C).

Together, the motif-level topology and the cell-type-grounded geodesics converge on a structural model of immune exclusion: the DFB shell, and specifically its interface sub-layer, separates a bulk stromal compartment holding trapped cytotoxic effectors from the malignant parenchyma. The model is independent of CD8^+^ abundance on two grounds — the 4 × 4 neighborhood Z matrices are abundance-normalized by construction, and the geodesic mixed-effects model treats sample-level CD8^+^ abundance as a random-intercept nuisance, isolating the within-sample effect of DFB coverage on CD8^+^ access. We emphasize that these analyses establish a strong, reproducible spatial association, not a perturbational test of causality.

#### Tumor-induced DFB expansion is associated with CD8^+^ T cell immune exclusion

Having shown that DFB architecture constrains CD8^+^ access within tumors, we asked whether the patient-to-patient degree of this exclusion tracks the tumor-induced expansion of the DFB compartment, and, critically, whether it does so beyond what stromal cell abundance alone would predict. We quantify expansion at the level of the DFB motif (the proportion of cells assigned the DFB barrier architecture), not CAF cell-type abundance: Delta DFB fraction therefore measures the growth of barrier-configured stroma (i.e., the reorganization of fibroblasts into a desmoplastic boundary, rather than simply more fibroblasts). Because baseline stromal composition varies across patients, we referenced each cancer section to its matched normal section and defined patient-specific DFB expansion as the change in DFB motif fraction between cancer and the projected normal baseline.

Across 16 paired colorectal cancer specimens, Delta DFB fraction ranged from −0.144 to 0.149 — some tumors contracted the DFB compartment relative to matched normal tissue while others markedly expanded it — and was strongly positively associated with the CD8^+^ core detour ratio, a graph-geodesic measure of how circuitous annotated CD8^+^ T-cell paths become when approaching the tumor core (Spearman ρ = 0.706, p = 0.0022; n = 16). Patients in the lowest detour-ratio quartile had a negative median Delta DFB fraction (−0.049), whereas those in the highest quartile had a clearly positive median (0.112): the tumors with the greatest DFB-motif expansion above their own normal baseline were those in which CD8^+^ T cells were forced into the most indirect routes to the core.

This association was not a restatement of stromal abundance. Normal-tissue DFB motif fraction was inversely associated with CD8^+^ detour ratio (Spearman ρ = −0.582, p = 0.0179), whereas the absolute cancer-tissue DFB motif fraction was positively associated (ρ = 0.729, p = 0.0013); the sign reversal indicates that constitutive stromal content in adjacent normal mucosa is not itself an immune-excluding state, and that exclusion tracks tumor-induced architectural remodeling above baseline rather than the amount of stroma present. Consistent with this, Delta DFB-motif expansion predicted CD8^+^ detour ratio after controlling for CAF cell-type abundance (partial Spearman ρ = 0.745, p = 0.0014), confirming that the effect reflects the configuration of the stromal compartment and not the number of fibroblasts.

The expansion was also linked to the specific architectural features expected of a barrier. Delta DFB fraction was positively associated with DFB boundary coverage around tumor regions (Spearman ρ = 0.685, p = 0.0034) and with the fraction of CD8-to-core paths crossing the DFB barrier (ρ = 0.576, p = 0.0194), with a weaker positive trend for DFB global continuity (ρ = 0.456, p = 0.0759). Together, these results support a model in which tumor-specific expansion of the DFB motif increases physical boundary formation around tumor regions, imposing longer and more tortuous migration routes for CD8^+^ T cells — an architectural, not merely compositional, determinant of CD8^+^ access.

#### Early-onset colorectal cancer amplifies the DFB-exclusion topology and yields an age-specific prognostic signature

We next asked whether the barrier-exclusion architecture differs between EOCRC and AOCRC and whether age-associated differences can be translated into a clinically interpretable signature. Differential expression between EOCRC and AOCRC within the DFB compartment identified a coherent activation program enriched for matrix-and-fibrosis biology (Figure 4A): the top gene-ontology terms in EOCRC-enriched DFB genes included supramolecular fiber organization, driven by COL1A1, COL1A2, COL3A1, COL5A1, COL6A3, MMP11, LUM, STMN1, DPYSL3, response to chemical, cellular response to organic substance, developmental process, and anatomical structure development. The leading-edge genes were dominated by fibrillar and basement-membrane collagens (COL1A1, COL1A2, COL3A1, COL4A1, COL4A2, COL5A1, COL6A3, COL17A1, COL18A1), matricellular proteins (SPARC, POSTN, LUM, IGFBP7, THBS1, VCAN), and matrix proteolysis (MMP11, CTSB), a coordinated EOCRC-specific upregulation of structural ECM biology within the same DFB motif identified in Figure 3. Beyond DFB-resident transcription, the global motif × motif interaction structure was reshaped in EOCRC relative to AOCRC along four pre-specified directional axes (Figure 4B). Tumor self-clustering was amplified in EOCRC (AO mean Z = 559.5 vs. EO 643.4, Δ +83.9, p = 0.0048), indicating tighter internal tumor parenchyma. DFB-Tumor exclusion was amplified in EOCRC (AO 173.9 vs. EO 212.1, Δ +38.2, p = 0.017), and CEH-Tumor exclusion was amplified in EOCRC (AO 48.5 vs. EO 64.6, Δ +16.1, p = 0.0031), confirming sharper tumor-stroma and tumor-effector segregation. CEH-DFB accumulation trended higher in EOCRC than AOCRC but did not reach significance (Δ +17.7, p = 0.22). Together, these axes describe a coherent EOCRC architectural phenotype: tighter tumor cores, a more pronounced DFB shell around them, and stronger trapping of cytotoxic effectors outside tumor parenchyma, exactly the structural-exclusion pattern motivated by Figure 3, intensified in early-onset disease. At the patient level, EOCRC tissues also carried a more abundant DFB compartment with thicker shell geometry (Figure 4C). The DFB size composite, a Z-summarized score combining DFB area, cell count, mean cell area, and median cell area, was higher in EOCRC than AOCRC (AO mean −0.282 vs. EO mean +0.282, Δ +0.564, p = 0.045), and the DFB thickness composite likewise separated EOCRC from AOCRC. These size and thickness measures are computed directly from motif maps and are independent of the interaction-Z analysis in Sup Fig. 3, providing an orthogonal confirmation that the EOCRC barrier-exclusion architecture is not only sharper at the interface but also larger and thicker in absolute terms.

**Figure 4.**
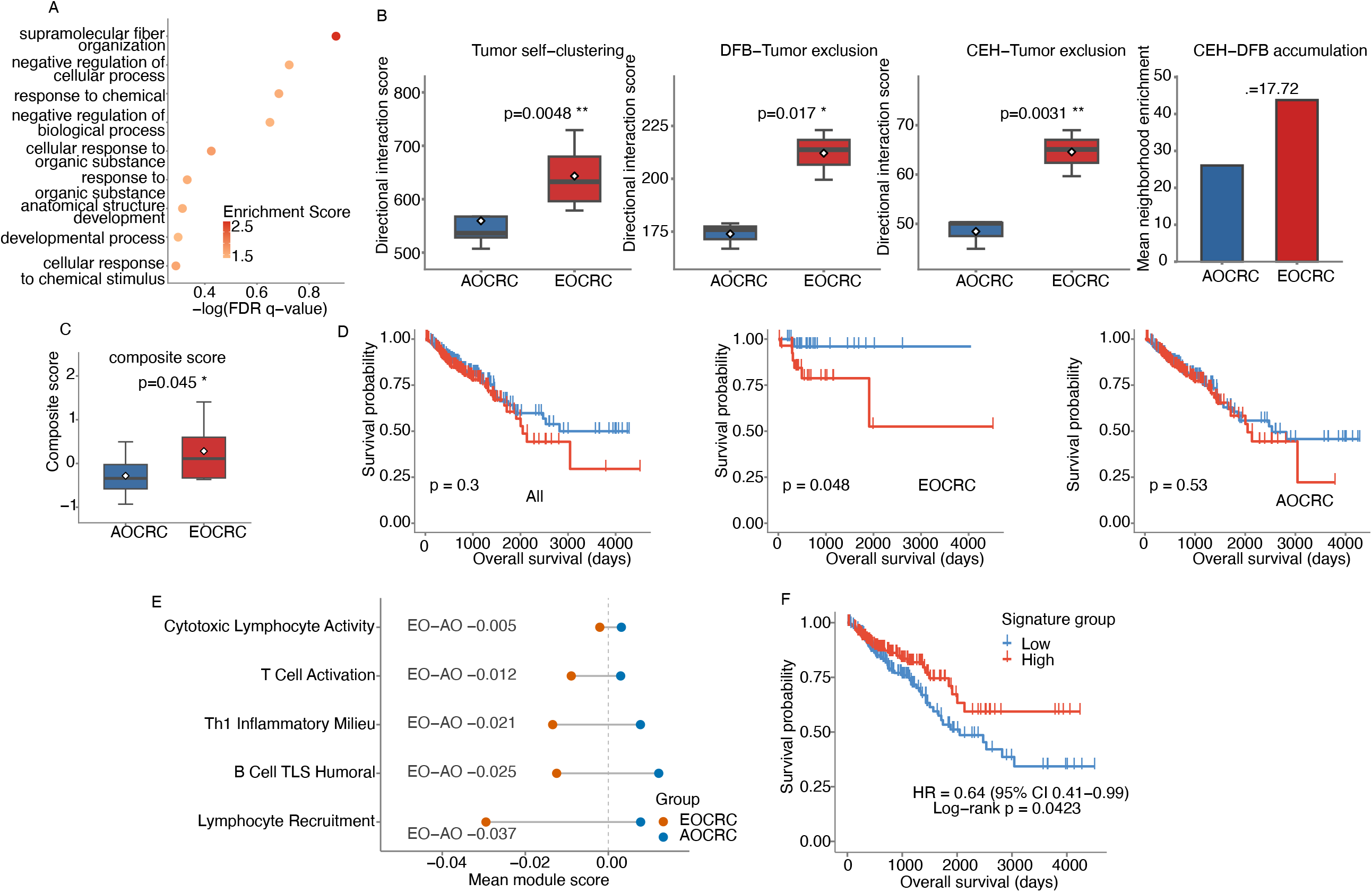
Early-onset CRC selectively amplifies the DFB-associated exclusion architecture and defines a DFB-associated biomarker. (A) Gene Ontology enrichment bubble plots for differentially expressed genes in EOCRC DFB. (B) Directional category boxplots: tumor self-clustering (P = 0.0048), DFB-tumor exclusion (P = 0.017), CEH-tumor exclusion (P = 0.0031), and CEH-DFB accumulation (P = 0.22). (C) DFB size composite per patient: AO −0.282 versus EO +0.282, one-sided permutation P = 0.045. (D) Kaplan-Meier analysis of TCGA COAD overall survival stratified by the EOCRC-DFB signature PC1 score. Patients were classified as signature-high or signature-low using the median PC1 score within each cohort. From right to left, curves are shown for all patients (n = 376), EOCRC patients <50 years (n = 57), and AOCRC patients ≥50 years (n = 319). The signature was not prognostic in the pooled cohort (HR = 1.26, 95% CI 0.82-1.92; log-rank P = 0.30) or in AOCRC (HR = 1.24, 95% CI 0.79-1.94; log-rank P = 0.34), but significantly stratified survival in EOCRC (log-rank P = 0.048), where signature-high patients showed shorter overall survival. (E) Immune-program scores within tumor-parenchymal cells, EOCRC versus AOCRC. For cells assigned to the four tumor-parenchymal motifs (SRN, PTC, DEC, IRN; motifs 4, 5, 7, 9), mean module scores are shown for five immune programs — cytotoxic lymphocyte activity, T-cell activation, Th1 inflammatory milieu, lymphocyte recruitment, and B-cell/TLS humoral activity — as a readout of immune infiltration into the tumor parenchyma. EOCRC tumor-parenchymal cells showed lower scores across all five programs than AOCRC, consistent with a more immune-excluded malignant compartment in early-onset disease. (F) Kaplan-Meier analysis of TCGA COAD overall survival stratified by the AOCRC-associated top-decile DEG signature. Patients were classified as signature-high or signature-low using the median signature score in all COAD patients (n = 376). Signature-high patients showed significantly improved overall survival compared with signature-low patients (HR = 0.64, 95% CI 0.41-0.99; log-rank P = 0.042).

We then asked whether the EOCRC-amplified DFB activation program is itself clinically meaningful, and whether its prognostic value is age-specific (Figure 4D). We defined an EOCRC-DFB signature from the top 10% differentially expressed genes between EOCRC and AOCRC DFB cells: MMP11, COL4A2, COL3A1, COL4A1, SPARC, COL1A2, CTSB, IGFBP7, and THBS1. This nine-gene panel spanning matrix proteolysis (MMP11, CTSB), fibrillar collagens (COL1A2, COL3A1), basement-membrane collagens (COL4A1, COL4A2), and matricellular proteins (SPARC, IGFBP7, THBS1). We computed a per-patient Z-score of this signature in the TCGA COAD cohort and stratified each cohort into high vs. low at the cohort median. In the pooled cohort (n = 376), the signature did not separate overall survival (HR = 1.26, 95% CI 0.82-1.92; log-rank p = 0.30). In the average-onset subset (≥50 years, n = 319), the signature was likewise non-significant (HR = 1.24, 95% CI 0.79-1.94; log-rank p = 0.34). In the early-onset subset (<50 years, n = 57), the same signature stratified survival with a substantially larger effect (p = 0.048), with early-onset patients in the signature-high group experiencing markedly shorter overall survival than those in the signature-low group. The wide confidence interval reflects the limited size of the early-onset TCGA COAD subset, but the age-stratified pattern — null in the pooled cohort, null in average-onset, significant only in early-onset. This analysis establishes the EOCRC-DFB signature as an age-specific prognostic readout that recapitulates the barrier-exclusion phenotype defined from spatial topology. Together, these results define a coherent EOCRC architectural phenotype with four orthogonal features: tighter tumor parenchyma, denser and thicker DFB shell, sharper DFB-Tumor and CEH-Tumor segregation, and a transcriptional DFB activation program enriched for fibrillar/basement-membrane collagens and matrix-remodeling proteases — and demonstrate that this phenotype translates into an age-specific prognostic signature in independent bulk-transcriptomic data. We note that the present cohort does not yet support full stage- or MSI-stratified inference at high power; analyses with stage and MSI as covariates, and replication in independent early-onset CRC cohorts, are listed as the highest-priority extensions.

Because EOCRC tumors showed an amplified DFB barrier, we next asked whether this architecture was accompanied by reduced immune infiltration into the tumor parenchyma. We reasoned that a denser DFB shell could limit the penetration of immune effectors into tumor-cell-rich regions, leaving the EOCRC malignant compartment comparatively immune-cold relative to AOCRC. To test this directly within the parenchyma, we restricted the analysis to cells assigned to the four tumor-parenchymal motifs (SRN, PTC, DEC, and IRN; motifs 4, 5, 7, and 9) and scored each of these cells for five immune programs — cytotoxic lymphocyte activity, T-cell activation, Th1 inflammatory milieu, lymphocyte recruitment, and B-cell/TLS humoral activity (Figure 4E). Scoring immune programs within tumor-motif cells provides a readout of how much immune signal reaches the tumor parenchyma, rather than the immune content of the tissue as a whole. Across all five programs, EOCRC tumor-parenchymal cells carried lower mean immune-program scores than their AOCRC counterparts. These results are consistent with an immune-exclusion model in EOCRC, in which the amplified DFB barrier is associated with reduced immune infiltration into the malignant compartment, whereas AOCRC tumors retain a more immune-hot parenchyma.

We then asked whether the AOCRC-enriched transcriptional program carried prognostic information in independent bulk-transcriptomic data. From the EOCRC versus AOCRC differential expression analysis, we selected the high-confidence top-decile AOCRC-associated DEGs and computed a per-patient signature score in TCGA COAD. Patients were stratified into signature-high and signature-low groups at the cohort median. In all TCGA COAD patients (n = 376), the AOCRC-associated DEG signature significantly separated overall survival, with signature-high patients showing improved survival relative to signature-low patients (HR = 0.64, 95% CI 0.41-0.99; log-rank p = 0.042; Figure 4F). Together with the EOCRC-DFB survival analysis, these data suggest opposing clinical associations for the two age-linked microenvironmental states: an EOCRC-amplified DFB/matrix program associated with immune exclusion and poorer outcome, and an AOCRC-enriched immune-hot program associated with more favorable survival.

#### H&E translation of the STORM motif framework yields a deployable prognostic biomarker in advanced-stage TCGA colorectal cancer

To broaden the clinical accessibility of the STORM framework beyond spatial omics, we asked whether motif-derived tissue classes could be detected directly from H&E whole-slide images. We registered Xenium motif maps to match H&E sections, extracted approximately 300,000 tiles labeled by majority motif macro-class (DFB, Immune, VBMS, or Tumor), and benchmarked six computer-vision backbones for tile-level motif classification (Figure 5A). ViT-B/16^29^ was the strongest model, achieving an accuracy of 0.82 and outperforming EfficientNet-B0^30^, DenseNet121^31^, and ResNet-18/34/50^32^.

**Figure 5.**
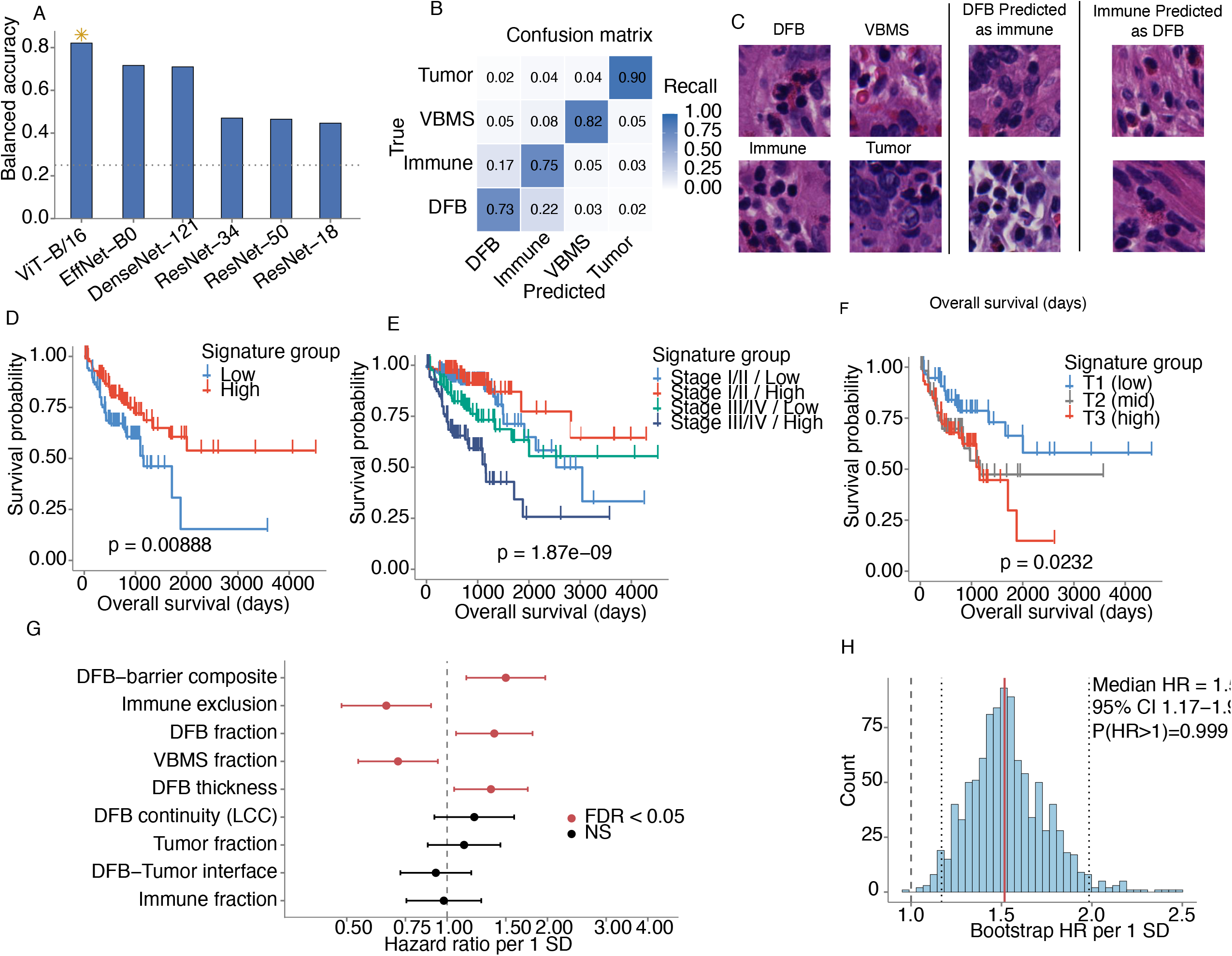
H&E translation of the STORM motif framework yields an advanced-stage prognostic biomarker in TCGA colorectal cancer. (A) Backbone comparison for tile-level motif macro-class classification across ViT-B/16, EfficientNet-B0, DenseNet121, and ResNet-18/34/50. (B) ViT-B/16 row-normalized four-class confusion matrix for DFB, Immune, VBMS, and Tumor motif macro-classes. (C) Representative TCGA whole-slide image overlaid with predicted motif macro-classes, with examples of DFB/Immune confused cases. (D) Kaplan-Meier curves for advanced-stage TCGA CRC stratified by immune-exclusion score. (E) Kaplan-Meier curves for advanced-stage TCGA CRC stratified by DFB-barrier composite tertiles. (F) Advanced-stage Cox forest plot (n = 177, events = 58), showing DFB-barrier composite (HR 1.50, q = 0.032), DFB fraction (HR 1.39, q = 0.035), DFB thickness (HR 1.35, q = 0.035), VBMS fraction (HR 0.71, q = 0.035), and immune-exclusion score (HR 0.66, q = 0.035). (G) Bootstrap stability analysis of the advanced-stage DFB-barrier composite Cox hazard ratio across 1,000 resamples, showing a consistently adverse association (mean HR = 1.54; P[HR > 1] = 0.999).

The ViT-B/16 confusion matrix showed robust recovery of the four H&E-translated motif macro-classes, with the largest residual confusion occurring between DFB and Immune tiles (Figure 5B). This error mode was biologically interpretable: spatial motif maps showed that DFB regions frequently surround or border immune motifs, producing H&E tiles in which fibroblastic barrier morphology and immune-rich tissue are closely interleaved. We therefore included representative DFB/Immune confused cases to illustrate that model uncertainty reflects the spatial organization of immune motifs embedded within or adjacent to DFB barriers, rather than arbitrary classification failure (Figure 5C).

We next applied the ViT motif classifier to TCGA colorectal cancer diagnostic slides and computed slide-level biomarkers, including DFB fraction, DFB thickness, DFB continuity, DFB-Tumor interface, an integrated DFB-barrier composite, immune-exclusion score, immune fraction, tumor fraction, and VBMS fraction. In the full TCGA cohort (n = 412, events = 90), these biomarkers showed directionally consistent effects but did not reach significance after FDR correction. In contrast, in the advanced-stage subset (stage III/IV; n = 177, events = 58), immune exclusion and the DFB-barrier composite showed clear survival-associated structure on Kaplan-Meier analysis (Figure 5D, E).

In advanced-stage Cox analysis, the DFB-barrier composite was the strongest prognostic feature (HR 1.50, 95% CI 1.14-1.97, p = 0.0036, FDR q = 0.032; Figure 5F). DFB fraction (HR 1.39, q = 0.035) and DFB thickness (HR 1.35, q = 0.035) were each associated with shorter overall survival, while VBMS fraction was protective (HR 0.71, q = 0.035) and immune exclusion retained a protective association under its score convention (HR 0.66, q = 0.035). A 1,000-iteration bootstrap further supported the stability of the DFB-barrier composite association, with HR > 1 in nearly all resamples (mean HR = 1.54; P[HR > 1] = 0.999; Figure 5G). Together, these results show that H&E-based motif translation can identify a biologically interpretable subset of advanced-stage CRC patients whose tumors are encased in a strong fibroblast barrier and whose prognosis is correspondingly worse.

#### Extension: LLM-assisted motif interpretation generalizes to independent datasets

To accelerate biological interpretation of unsupervised STORM motifs, we developed STORM-Interpret, an agentic LLM pipeline that annotates motifs from their differential-expression signatures after unsupervised discovery, preserving statistical independence between motif definition and interpretation (Sup File 2 and Sup Fig 4). Applied to two independent Xenium datasets, which are rhabdomyosarcoma (10 motifs) and lung cancer (5 motifs). The pipeline generated structured annotations with gene-citation validation for 14 of 15 motifs. Notably, both datasets contained a fibroblast- and ECM-rich stromal motif annotated as a desmoplastic barrier-like structure, consistent with the DFB architecture identified in our colorectal cancer cohort and suggesting that DFB-like spatial architectures may recur across tumor types. These exploratory observations require validation with dedicated spatial analyses but indicate that LLM-assisted interpretation can accelerate first-pass motif annotation without biasing motif discovery.

## Discussion

In this study, we developed STORM, an unsupervised graph-attention variational autoencoder framework for discovering recurrent multicellular spatial motifs from single-cell-resolution spatial transcriptomic data. Applied to paired colorectal cancer Xenium sections from early-onset and average-onset patients, STORM identified ten recurrent spatial motifs that organized into tumor-parenchymal, stromal, and immune macro-compartments without using cell-type labels, manual tissue annotations, or predefined tumor-stroma-immune compartments. This annotation-free organization suggests that colorectal tumors contain reproducible spatial rules that can be recovered directly from the joint structure of molecular states and local tissue topology.

The central finding of this work is the identification of a Desmoplastic Fibrotic Barrier (DFB) motif as a recurrent stromal boundary architecture associated with restricted cytotoxic T-cell access to tumor cores. Importantly, DFB is an architectural label, a motif assigned to cells by their neighborhood context and should not be read as a fibroblast-rich or ECM-rich cell type. STORM discovered DFB as a single motif. The interface and non-interface labels are not separate motifs. We defined them downstream by splitting DFB-motif cells on tumor adjacency. Using this split, our analyses support a layered spatial model. Interface DFB cells directly contact tumor parenchyma. Non-interface DFB cells form a surrounding bulk stromal compartment. This bulk compartment co-localizes with the cytotoxic effector hub (CEH) motif. Crucially, when we grounded this motif-level topology in the independent cell-type annotation, the annotated CD8^+^ T cells themselves were not absent from the tissue but accumulated within this bulk stromal compartment while remaining spatially displaced from malignant cores. This convergence of an architectural motif and an independent cell-type annotation refines the conventional concept of immune exclusion: exclusion may arise not only from reduced immune infiltration, but also from the geometric organization of stromal barriers that separate cytotoxic cells from tumor cores.

A major conceptual contribution of this study is the shift from composition-based to topology-based interpretation of the tumor microenvironment. Many spatial and single-cell analyses quantify the abundance, proximity, or co-localization of CAFs, tumor cells, and immune cells. However, abundance and Euclidean proximity alone do not determine whether cytotoxic cells can physically access malignant regions^33^. By modeling tissue as a cell-level spatial graph, we quantified CD8^+^ T-cell access using graph-geodesic distances, barrier-crossing fractions, and DFB-weighted detour ratios. These analyses showed that stronger DFB boundary coverage was associated with a greater probability that CD8^+^ paths must traverse DFB and with increased geodesic cost to reach tumor cores. Thus, the DFB motif acts as a spatial-topological constraint on immune access rather than merely a compositional stromal feature.

The abundance-independent nature of this result is important. The neighborhood-enrichment framework compares observed motif adjacencies to a random-permutation null that preserves motif counts, allowing DFB-tumor, CEH-tumor, and CEH-DFB relationships to be interpreted beyond simple differences in cell abundance. Similarly, the tile-level mixed-effects models evaluate within-sample variation in DFB boundary coverage while accounting for sample-level heterogeneity. Together, these analyses support a model in which the spatial arrangement of DFB, rather than CD8^+^ abundance alone, contributes to impaired tumor-core access. We nevertheless emphasize that these data establish a strong and reproducible spatial association, not direct perturbational causality.

Our results also suggest that early-onset colorectal cancer may differ from average-onset disease at the level of tissue architecture. EOCRC tissues showed a coordinated amplification of the DFB-exclusion topology, including stronger DFB-associated ECM activation, increased DFB size and thickness, sharper DFB-tumor segregation, and stronger CEH-tumor exclusion. These findings suggest that EOCRC may not only differ in genetic, molecular, or clinical features, but may also exhibit a distinct stromal architecture that promotes immune exclusion. The EOCRC-enriched DFB transcriptional program, including collagens, matrix-remodeling genes, and matricellular proteins, further supports the idea that early-onset tumors may activate a more pronounced fibroblast barrier state. The association between the EOCRC-DFB signature and poorer survival specifically in younger TCGA-COAD patients provides an independent transcriptomic readout of this age-associated barrier phenotype, although this finding requires validation in larger early-onset cohorts.

The H&E translation analysis extends the biological findings toward clinical applicability. Although high-resolution spatial transcriptomics is powerful, it remains costly and is not yet routinely available for large clinical cohorts. By training a whole-slide image classifier to recognize motif-derived macro-classes from H&E tiles, we asked whether spatial motifs learned from Xenium could be approximated from routine pathology images. The ViT-based classifier achieved sufficient performance to generate slide-level motif maps, from which DFB-centered biomarkers were computed in TCGA colorectal cancer slides. In advanced-stage disease, the image-derived DFB barrier composite was associated with shorter overall survival and remained stable under bootstrap resampling. This result suggests that spatial motifs discovered from molecular tissue maps can be translated into scalable pathology biomarkers that capture clinically relevant stromal architecture. These findings have several broader implications. First, they support the view that tumor progression and immune escape are shaped not only by cellular states, but also by the spatial architectures that constrain how cells interact. Second, they suggest that fibroblast biology should be interpreted in relation to tissue geometry: the same CAF-associated programs may have different functional consequences depending on whether they form diffuse stroma, perivascular scaffolds, or circumferential tumor barriers^34^. Third, they motivate the development of topology-aware biomarkers that quantify structural access, barrier continuity, and immune displacement rather than relying only on cell fractions or marker intensity. Such biomarkers may be particularly relevant in advanced-stage colorectal cancer, where stromal organization may influence whether endogenous or therapy-induced immune responses can reach malignant regions.

Several limitations should be noted. First, the spatial transcriptomic cohort is modest in size and derived from a single institutional setting, with nine EOCRC and seven AOCRC patients. Although STORM motifs were recurrent across samples and multiple analyses supported the DFB-exclusion model, larger independent cohorts will be needed to validate the robustness and generalizability of the EOCRC-specific architectural phenotype. Second, Xenium measures a targeted gene panel rather than the whole transcriptome. This design enables high-quality spatial profiling of selected CRC and TME programs, but may miss additional immune, stromal, or metabolic states relevant to tumor architecture. Third, the current study is observational. The geodesic analyses demonstrate that DFB boundary organization is associated with impaired CD8^+^ access, but perturbational experiments are required to determine whether disrupting DFB-associated ECM programs restores tumor-core immune infiltration. Fourth, H&E-based motif classification remains imperfect, particularly for morphologically similar stromal and immune classes. Although slide-level aggregation recovered prognostic signal, improved pathology models, larger training datasets, and external validation will be needed before clinical deployment. Finally, TCGA survival analyses are retrospective and may be influenced by treatment, MSI status, CMS subtype, slide quality, and other clinical covariates that were not fully modeled in the current analysis. Future work should validate the DFB barrier model using orthogonal spatial proteomic and experimental systems. Multiplex immunofluorescence, CODEX, or imaging mass cytometry could directly test the predicted layered organization of interface DFB, bulk DFB, and CD8^+^ effector cells at the protein level. Perturbational models targeting DFB-associated ECM and matrix-remodeling genes, such as collagen programs, MMP11, SPARC, THBS1, or related CAF pathways, could determine whether weakening the DFB barrier increases CD8^+^ tumor-core access. Larger EOCRC cohorts with matched clinical, molecular, MSI, and treatment information will also be important for testing whether the DFB-exclusion phenotype explains aggressive behavior in specific early-onset subgroups. More broadly, applying STORM to other solid tumors may reveal whether fibroblast-mediated geodesic exclusion represents a general architectural principle of immune evasion.

In summary, STORM provides a topology-aware, annotation-free framework for discovering recurrent spatial motifs across patients. In colorectal cancer, this approach identified a desmoplastic fibrotic barrier architecture that spatially separates cytotoxic effectors from tumor cores, is selectively amplified in early-onset disease, and can be translated into H&E-derived prognostic biomarkers in advanced-stage colorectal cancer. These findings support a model in which immune exclusion is not only a matter of cell abundance or immune activation state, but also a consequence of recurrent tissue architectures that constrain physical access within the tumor microenvironment.

## Materials and Methods

### Technical overview of STORM

#### Motivation and design rationale

A recurrent spatial motif is, by our definition, a multicellular configuration determined jointly by the molecular states of cells and their spatial arrangement, and reproducible across patients. This definition imposes three requirements on any model intended to discover such motifs de novo. First, the representation of each cell must encode not only its own transcriptional profile but also the composition and geometry of its local tissue neighborhood, so that two transcriptionally similar cells occupying different architectural contexts are assigned distinct representations. Second, because no ground-truth motif labels exist and we wish to avoid imposing predefined cell-type annotations or compartment definitions, learning must be fully unsupervised. Third, the representation must be comparable across patients so that motifs recur as reproducible units rather than as sample-specific clusters. STORM addresses these requirements with a graph-attention variational graph autoencoder (GAT-VGAE) that (i) propagates information along a spatial proximity graph so that each cell’s embedding is contextualized by its neighbors, (ii) is trained by reconstructing both expression and local graph structure under a variational prior, and (iii) is fit jointly across all patient graphs to yield a single shared embedding space in which motifs are defined by clustering. Joint training and pooled clustering impose cross-patient comparability by construction; whether the resulting vocabulary elements actually *recur* — i.e., are molecularly reproducible across patients — is an empirical question evaluated separately (recurrence AUC, Fig. 1E), not an assumption built into the model.

#### Notation and spatial graph construction

Suppose we have spatial transcriptomics samples from *p* = 1, ···, *P* patients with measurements on *G* genes. After data preprocessing (cell segmentation and quality control), each patient *p* contributes *C* _*p*_ cells with gene expressions 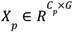 and spatial coordinates 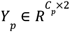. From *Y*_*p*_, we construct a weighted k-NN graph Ψ_*p*_ = (*V*_*p*_, *E*_*p*_, *w* _*p*_), where each node in *V*_*p*_ = {1, ···, *C*_*p*_} is a cell, each edge in *E*_*p*_ ⊂ *V*_*p*_ *x V*_*p*_ encodes local spatial adjacency, and *w* _*p*_: *E* → R_*p*_ assigns each edge a weight. The associated adjacency matrix *A*_*p*_ has *A*_*p*_ (*i, j)* = 1 if (*i, j)* ∈ *E*_*p*_ and 0 otherwise. Each node *i* carries as its feature vector the cells’s scaled expression profile *x*_*i*_ ∈ *R*^*G*^.

Throughout we set *K* = 10 and

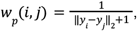

where *y*_*i*_, *y*_*j*_ are the *i*th and *j*th rows of *Y*_*p*_, and ‖ · ‖_2_is the Euclidean norm. The + 1 in the denominator bounds the weight and avoids division by zero at coincident coordinates; the monotone decay in distance encourages closer neighbouring cells to contribute more strongly during graph message passing, embedding the assumption that spatial proximity reflects stronger local tissue interaction.

#### Generative model

Assuming patients are independent, we drop the subscript *p* for clarity. Following the VGAE^35^ framework, we associate each cell *i* ∈ *V* with a latent embedding *z*_*i*_ ∈ *R*^*d*^ with *d* = 20 and collect *Z* = {*z* _1_, ···, *z*_*C*_}. The generative model factorizes the joint likelihood of expression and graph structure as

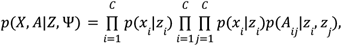

with a Gaussian feature decoder and a Bernoulli (inner-product) structural decoder:

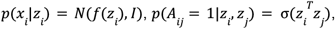

where *f*(·) is parameterized by a deep neural network, and σ(·) is the logistic sigmoid. The two decoders play complementary and deliberately chosen roles: the feature decoder forces the embedding to retain the cell’s molecular identity, while the structural decoder forces embeddings of spatially adjacent cells to be predictive of their adjacency, i.e. to lie close in the latent geometry. It is precisely this structural objective that makes each *z* _*i*_ a representation of the cell in its niche rather than of the cell in isolation, the embedding must reconstruct who a cell’s neighbours are, so cells sharing a recurrent neighbourhood composition are pulled together regardless of fine transcriptional differences, while transcriptionally similar cells in different neighbourhoods are pushed apart. We place an a-priori-independent standard normal prior 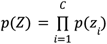 with *p*(*z* _*i*_) = *N*(0, *I)*, following the standard VAE setup^36^.

#### Inference model

We adopt a mean-field variational posterior

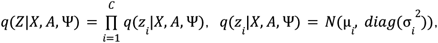

whose parameters μ _*i*_ and 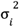 are produced by a two-layer GATv2 encoder^37^. Unlike the GCN encoder of the original VGAE, GATv2 learns anisotropic attention coefficients over each cell’s neighbourhood, allowing the model to weight neighbours unequally, appropriate for tissue, where a cell’s architectural context is dominated by a subset of its neighbours. The scalar edge weights *w* are supplied as edge features and incorporated into the attention logits, so that physical proximity modulates the learned attention in addition to the message magnitude. Because the same encoder weights are shared across all patient graphs, the resulting embeddings inhabit a common latent space and are directly comparable across patients, the property that lets motifs recur as cross-sample units.

#### Training objective

Model parameters are fit by maximizing the associated evidence lower bound (ELBO). Because patients are independent, the cohort ELBO is the sum of per-patient ELBOs. Following the β-VAE^38^ formulation, each per-patient ELBO is

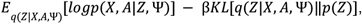

Where β ≥ 0 controls the strength of the prior regularization and the reconstruction term decomposes into the expression and structural likelihoods defined above. Throughout we set β = 0. 005 to prevent posterior collapse as downstream motif discovery depends on the latent embeddings being discriminative. Expectations are estimated with the reparameterization trick. Due to the large number of cells, evaluating the full structural likelihood is infeasible.

#### Spatial motif construction

After convergence, we discard the stochastic component and take the posterior mean μ _*i*_ as the point embedding of cell *i*, giving a single^d^-dimensional representation per cell in the shared latent space across all *P* patients. Because these embeddings jointly encode molecular state and local graph context, clustering them groups cells by recurrent neighbourhood architecture rather than by transcriptional identity alone. We pool the embeddings of all cells from all sections and partition them with mini-batch k-means into *M* clusters; each cluster defines one spatial motif, and every cell receives a motif label corresponding to its cluster assignment. We selected *M* = 10. Motif labels were then mapped back onto the original spatial coordinates of each section for downstream biological interpretation.

#### Xenium data generation

Patients with either EO-CRC or AO-CRC were selected from our institutional cohort at UTSW MC after obtaining informed consent (IRB STU 21-0161). Patients with familial or hereditary CRC, MSI status, and/or history of inflammatory bowel disease were excluded. We also excluded subjects without an available matched normal tissue specimen. H&E slides were created for both the tumor and matched normal tissue. The presence of tissue of interest, for example cancer, was confirmed and marked by a board-certified gastrointestinal pathologist (ZC).

In situ spatial transcriptomic testing was conducted on surgical specimens that were fixed in formalin and embedded in paraffin using 10x Genomics’ Xenium platform. Prior to processing the samples, RNA quality was verified. Briefly, three 10um Formalin-fixed, Paraffin-embedded (FFPE) tissue curls per sample were evaluated for RNA quality, excluding samples with DV200 <30%. Then, the tissue was sectioned onto the Xenium slides following the Xenium In Situ for FFPE - Tissue Preparation Guide (CG000578, Rev C). Briefly, 5um sections were mounted onto 10x Genomics Xenium slides and adhered via baking at 42°C for 3 hours followed by overnight incubation at 37°C. The samples then were deparaffinated through sequential immersion in xylene and graded ethanol, followed by rehydration. Tissue quality and morphology were validated using Hematoxylin and Eosin (H&E) staining and DAPI assessment.

A pre-designed, off-the-shelf Human Colorectal Gene Set of 325 genes from 10x genomics and a unique 50-gene set to subclassify CAFs into general fibroblasts, inflammatory CAF (iCAF), myofibroplastic CAF (myCAF), and antigen-presenting CAFs (apCAF) were used for in-situ transcriptome mapping^39^. The CAF subclassification gene list was extracted from a Pan-Cancer CAF transcriptomic atlas^40^.

FFPE Xenium sample preparation and in situ gene expression were conducted following the 10x Genomics protocol (CG000578; CG000582). Briefly, formalin-fixed paraffin-embedded (FFPE) human colon cancer biopsy tissue was sectioned at 5 µm using a microtome and mounted onto Xenium slides following RNA quality assessment. Sections were dried at room temperature for 10 minutes and baked at 42°C for 3 hours to ensure tissue adhesion. Deparaffinization was performed using fresh xylene, followed by rehydration through a graded ethanol series (100%, 100%, 96%, 96%, 70%) and equilibration in nuclease-free water. Tissue sections then underwent decrosslinking to reverse formaldehyde-induced crosslinks and improve RNA accessibility while preserving tissue morphology.

Custom-designed Xenium probe panels (Xenium Design ID AU3TC8) targeting genes associated with colorectal cancer biology and tumor microenvironment characterization were hybridized to tissue sections overnight (16-24 hours) using a thermocycler-based incubation system. On the second day, unbound probes were removed by post-hybridization washing. Hybridized probe pairs were then ligated to generate circular DNA probes on target RNA molecules. Circularized probes underwent rolling-circle amplification to produce amplified barcode concatemers for downstream imaging detection.

Following amplification, tissue autofluorescence was reduced using Xenium autofluorescence quenching reagents, and cell nuclei were counterstained with DAPI nuclei staining buffer. Processed slides were loaded onto the Xenium Analyzer for automated cyclic fluorescence imaging, transcript decoding, and spatial transcriptomic data acquisition at subcellular resolution (CG000584). Xenium spatial transcriptomics assay was performed in the Microbiome Core at UT Southwestern Medical Center.

#### Data process and cell type annotation

For Xenium data preprocessing, we first manually registered the histology image to the Xenium spatial coordinate system to align tissue morphology with cell-level transcriptomic measurements. The registration quality was visually inspected to ensure accurate correspondence between image regions and Xenium cell coordinates. After image registration, we processed the Xenium count matrix using a standard scanpy^41^ workflow. Raw counts were loaded into an AnnData object together with cell metadata and spatial coordinates. Low-quality cells and rarely detected genes were filtered out based on total transcript counts and the number of detected genes. The filtered data were then library-size normalized, log-transformed using log1p, and used for downstream analysis. The expression matrix was normalized to a constant library size per cell, log-transformed, and scaled before dimensionality reduction. Principal-component analysis was performed, followed by construction of a cosine-distance k-nearest-neighbor graph and Leiden clustering^42^. Cell clusters were manually annotated using canonical marker-gene sets curated from prior literature and restricted to genes included in the Xenium panel. Tumor and immune populations were identified using markers. Annotation confidence was evaluated based on concordance among multiple markers, cluster-level differential-expression analysis, DotPlot visualization, spatial localization, and cellular morphology. Clusters showing heterogeneous or overlapping marker profiles were iteratively subclustered at resolutions selected according to the expected complexity of the corresponding population. Cells or subclusters lacking a coherent marker profile, exhibiting poor segmentation, or displaying incompatible marker combinations suggestive of merged cells were excluded from downstream analysis. The resulting annotations represented well-defined tumor, immune, vascular, and stromal cell populations at single-cell spatial resolution.

#### Benchmark

The benchmark comprised 2,794,520 cells from 16 colorectal cancer Xenium samples, including nine early-onset and seven average-onset cases, measured with a shared 375-gene panel. Downstream scores were calculated from persisted embeddings and motif labels without refitting the representation models during metric computation. STAGATE and SpaceFlow were fitted separately within each sample. To accommodate dense graph construction at this single-cell scale, GraphST, SEDR, and SpaGCN were fitted within contiguous spatial tiles containing at most 25,000 cells; the GraphST tile limit was reduced to 15,000 cells for samples that did not complete at the larger setting. Tile-level embeddings were restored to the original cell order, and the method wrappers standardized latent dimensions within each sample or tile. For each comparator, the pooled cohort embedding was clustered into K=10 (motifs using:batch size, 4,096 and random seed, 42) The STORM production labels were generated from the pooled STORM embedding with the same K, batch size, initialization setting, and random seed. Therefore, Figure 1B evaluates the consistency of globally defined motif labels within this cohort rather than independently discovered patient-specific labels. Comparator settings were fixed before scoring and were not exhaustively optimized.

Marker-program decodability. The prediction targets were 16 scanpy score_genes module scores calculated from predefined cell-type marker sets on the normalized 375-gene expression panel. Let h_pi_ ∈ ℝ^d^ denote the method-specific embedding of cell i from patient p, and let y_pij_ denote marker-module score j. Within each training fold, ridge-regression coefficients were estimated as:

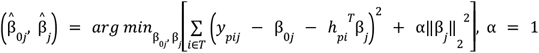

Models were evaluated using shuffled five-fold cross-validation, so every cell received a prediction from a model that had not been fitted to that cell. For patient p, method m, and module j, out-of-fold prediction performance was:

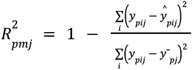

The patient-level metric was the mean across the J_p_ nonconstant module scores:

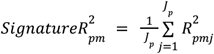

Thus, a higher signature R^2^ indicates greater linear predictability of the predefined marker-module scores from the embedding. Within each patient and method, cells with nonfinite embedding values were excluded and the remaining cells were randomly subsampled, when necessary, to a maximum of 50,000 cells (seed, 42). STORM embeddings were joined to the benchmark cells by patient and spatial coordinates rounded to two decimal places; 97.57% of cells had finite matched STORM embeddings and were eligible for Figure 1D. Because finite-value masks were applied separately by method, eligible cell sets could differ slightly between methods before subsampling. The module scores were calculated from the same expression panel available to the representation methods and had been generated during the cell-typing workflow; signature R^2^ was therefore interpreted only as marker-program decodability, not as independent biological ground truth. Cross-patient motif recurrence. Log-normalized expression was standardized gene-wise across the pooled cohort. Let x_pi_ ∈ ℝ^375^ denote the standardized expression vector of cell i from patient p, and let *ℓ*_pi_ ∈ {1,…,K} be its motif assignment. For each patient-motif combination containing at least 50 cells, the motif profile was:

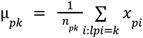

For a held-out patient p, the reference profile for motif k was the mean profile among the remaining patients in which that motif was present:

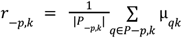

Cosine similarity between held-out motif k and reference motif l was:

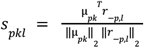

The recurrence AUC for patient p was the proportion of matched-versus-nonmatched comparisons in which the matched similarity was greater:

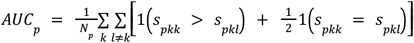

Here, N_p_ is the number of valid matched-versus-nonmatched comparisons. This statistic is the probability that a held-out motif profile has greater cosine similarity to the reference profile with the same global label than to a reference profile with a different label. Under no preferential matching, the expected AUC is 0.5; an AUC of 1.0 indicates that every valid matched comparison exceeds its nonmatching counterpart. A matched null distribution was also generated using 2,000 independent per-patient derangements of motif indices. Figure 1B and the STORM-versus-peer tests used the observed patient-level AUCs rather than null-corrected differences. Because motif clustering and gene standardization were performed using the pooled cohort, this metric measures within-cohort consistency and should not be interpreted as fully external out-of-sample performance.

The patient was the unit of analysis. For each method and metric, the plotted value was the arithmetic mean of the patient-level scores:

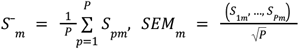

For each STORM-versus-peer comparison, the paired difference was *d*_p_ = *S*_p,STORM_ − *S*_p,peer_, because higher values were favorable for both Figure 1 metrics. A one-sided paired Wilcoxon signed-rank test evaluated the alternative that the patient-level differences were greater than zero. The paired effect size was

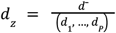

Uncertainty in the mean paired difference was estimated using 10,000 patient-level bootstrap resamples, with the 2.5th and 97.5th percentiles defining the 95% confidence interval. Within each metric, *P* values were Holm-adjusted^43^ across all *M* = 5 non-STORM methods in the complete benchmark. If *p*_(1)_ ≤ … ≤ *p*_(M)_ are the ordered raw *P* values, the adjusted value for rank *i* was

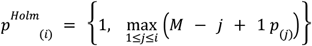

#### Spatial interaction matrix

Spatial neighborhood graph. For each sample, a spatial neighborhood graph was constructed from cell centroid coordinates using a k-nearest-neighbor scheme (squidpy.gr.spatial_neighbors, coord_type=“generic”, k=10). Let *G* = (*V, E)* denote the resulting undirected graph, where each node *v* ∈ *V* is a cell annotated with a spatial-motif label *l* (*v*) ∈ 0, 1, … 9, and each edge (*u, v)* ∈ *E* indicates spatial adjacency. The adjacency matrix *A* was symmetrized and its diagonal set to zero.

Observed interaction counts. For each ordered pair of motifs (*i, j)*, we counted the number of edges connecting a cell of motif *i* to a cell of motif *j*:

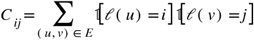

with each undirected edge counted once. Off-diagonal entries *C*_*ij*_ quantify heterotypic adjacency between motifs *i* and *j*, whereas diagonal entries *C*_*ij*_ quantify homotypic (self-)aggregation within a motif.

Permutation null model. Because *C*_*ij*_ depends on the marginal abundance of each motif, we assessed enrichment against a degree- and composition-preserving null. The motif labels were randomly permuted across all nodes while holding the graph topology *A* and the per-motif label counts fixed. For each of *b* = 300 permutations *b*, we recomputed the interaction matrix *C*^(*b)*^ and estimated the null mean and standard deviation for every pair:

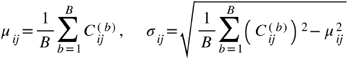

Enrichment statistics. The interaction strength for each motif pair was expressed as a z-score of the observed count relative to the null distribution:

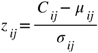

Positive *z*_*ij*_ indicates that motifs *i* and *j* are co-localized (spatial attraction) more frequently than expected by chance, whereas negative values indicate spatial avoidance/segregation. A one-sided empirical p-value for enrichment was computed as

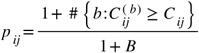

The resulting 10 *x* 10 z-score matrix defines the motif interaction map for each sample and was visualized as a heatmap. Per-sample matrices were retained for downstream aggregation across samples.

#### Graph Topology analysis

Spatial graph. For each sample we model tissue as a graph *G* = (*V, E)* whose nodes are cells and whose edges connect spatial neighbors. Edges are obtained from a *K*-nearest-neighbor graph on the cell centroids, symmetrized so that *A* = *A*^T^, with self-loops removed. All “distances” below are graph geodesic distances *d*_*G*_(*u, v*) i.e., the minimum number of edges (hops) on a shortest path between two cells, computed by breadth-first search (BFS)^44^ rather than Euclidean distance, so that the metrics follow tissue connectivity rather than straight-line space.

Each cell carries a spatial-motif label. We define two node sets: the DFB set *D* (DFB) and the tumor set *T* (Tumor motifs).

Tiling. To control for edge effects and within-sample heterogeneity, each sample is partitioned into a 4 *x* 4 grid of tiles. Every tile is grown by a 250 µm buffer (“halo”); the graph and all geodesic computations use the buffered patch, but every reported quantity is restricted to the inner (un-buffered) cells. Tiles failing minimum-count QC (cells, tumor cells, DFB cells, interface-DFB cells, core cells) are skipped. Per-tile values are then aggregated to the sample level.

Tumor core and boundary. A tumor cell is a boundary cell if it has at least one non-tumor neighbor:

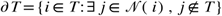

Using ∂ *T* as multi-source seeds, we compute the hop distance of every tumor cell to the tumor margin, d _∂_*(i)=d*_*G*_*(i ∂T)*, and define the tumor core as cells lying deep inside the mass:

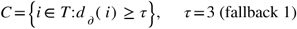

Interface DFB. The DFB cells that directly abut tumor define the shell candidate set:

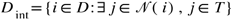

Angular coverage and orientation entropy. Let 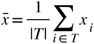 be the tumor centroid. Each interface-DFB cell is assigned a polar angle relative to the centroid:

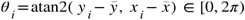

and angles are histogrammed into *B* = 120 bins (5° each) with occupancy counts *n*_*b*_ Angular coverage is the fraction of bins that are occupied (>1 cell):

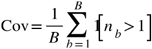

which measures how completely DFB encircles the tumor (1 = full wrap). The angular distribution 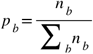 gives a normalized orientation entropy

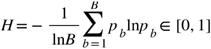

with *H* → 1 for uniform encirclement and low *H* for one-sided accumulation.

Must-cross fraction (geodesic encirclement test). This quantifies whether DFB topologically separates the tumor core from its surroundings. Let *O* be the non-tumor “outside” cells in the tile. We compute, on the full graph, the set of outside cells geodesically reachable from the core *C,R* _full_ ={*0* ∈O:*d*_*G*_*(o,C)* < ∞} We then delete all DFB nodes (except any in the core) from the graph and recompute reachability^R^_block_. A path is forced through the DFB shell if it exists in the full graph but not after DFB removal:

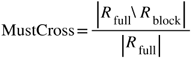

A value near 1 means almost every route between the tumor core and the outside is intercepted by DFB, i.e., a continuous enclosing barrier.

Robustness. All topology metrics are recomputed over a sweep of graph connectivity (*k* ∈{6,10,14,18}) to confirm that conclusions are not artifacts of a particular neighborhood size.

Detour ratio. On the same symmetrized *K*-NN cell graph, we treat DFB cells as a soft barrier set *B* and ask how much that barrier lengthens the path an external CD8^+^ T cell would have to travel to reach the tumor core. The source set is the tumor core *C*. The agents are the external CD8^+^T cells,*={i:*CD8, *i*. ∉*T*,*I* ∈ inner tile} All distances are graph geodesics; tiling and inner/halo handling are identical to the shell-geometry analysis.

Unweighted geodesic distance. Ignoring the barrier, the hop distance from the core to each cell is obtained by multi-source BFS from *C*:

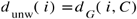

Barrier-weighted geodesic distance. We assign each node a traversal cost that is elevated inside DFB,

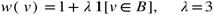

so stepping onto a DFB cell costs 1+ λ = 4 versus 1 elsewhere. The weighted distance is the minimum-cost path from the core, computed by multi-source Dijkstra^45^:

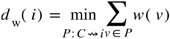

This lets a path either pay to pass through DFB or route around it, whichever is cheaper. For each external CD8^+^ cell with a finite, positive distance, the detour ratio is the inflation of path cost caused by the barrier,

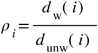

reported per tile as the mean and median over eligible CD8^+^ cells:

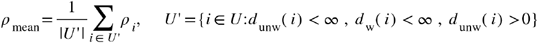

Interpretation. ρ_*i*_*=*1 means the cheapest route from the tumor core to that CD8^+^ cell never crosses DFB — the immune cell has an unobstructed approach. ρ _i_ > 1 means every route is either forced through DFB (paying the per-node penalty) or must take a longer barrier-avoiding detour; larger values indicate that DFB topologically insulates the tumor core from immune cells. Tiles are aggregated to the sample level, and the ratio is read alongside the complementary barrier-cross fraction (the fraction of CD8^+^ cells whose shortest path to the core steps onto at least one DFB node) to separate “must pass through DFB” from “must go around it.”

#### Tumor-induced DFB expansion predicts CD8^+^ immune exclusion analysis

Two separate STORM (Spatial Topology analysis of Recurrent Motifs) models were independently trained on the 16 cancer sections and 16 normal sections respectively. Each model was configured with K = 10 spatial motifs, identified by k-means clustering of the latent embeddings. Because the cancer and normal STORM models were trained independently, their motif spaces are not directly comparable. To project normal motif fractions into the cancer motif coordinate system, we computed a 10 × 10 Jaccard similarity matrix J between the top-50 marker gene sets of each cancer motif and each normal motif. The best-matching normal motif for cancer motif DFB was normal motif 0 (Jaccard = 0.351). The Jaccard-weighted projection was defined as:

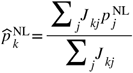

where 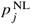 is the fraction of cells assigned to normal motif *j* in the patient’s normal tissue section, and 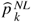 is the corresponding projected weight in cancer motif space *k*.

For each patient, the per-cancer-motif delta was computed as:

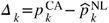

where 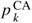 is the fraction of cancer cells assigned to cancer motif *k*. Δ DFB refers specifically to Δ_0_ (cancer motif 0). A positive value indicates that the DFB niche is proportionally more abundant in the patient’s cancer tissue than predicted by their own normal tissue baseline.

#### Pathology Translation

Training-patch construction and motif labels. To learn an image-only readout of STORM spatial tissue motifs, we constructed a labelled H&E patch dataset from Xenium samples with co-registered H&E images. For each segmented cell with an upstream motif assignment and a matched H&E coordinate, we extracted one RGB patch centered on the cell and assigned the patch the corresponding spatial-motif label. Original motif identities were collapsed into four interpretable macro-classes for supervised learning: DFB, Immune, VBMS, and Tumor. Patch-label matching was performed through a manifest linking each per-cell H&E patch to its motif annotation.

Backbone comparison and model selection. We trained a four-class patch-level classifier and compared six ImageNet-pretrained backbones under the same training and evaluation framework: ResNet-18, ResNet-34, ResNet-50, DenseNet-121, EfficientNet-B0, and ViT-B/16. Held-out test data were split at the sample level, ensuring that no cells from a test slide appeared during training. Remaining patches were divided into training and validation sets using a stratified 85/15 split. Class imbalance was handled with class-balanced sampling during training and inverse-frequency class weights in the loss. Input patches were center-cropped around the labelled cell, resized to 224 × 224, normalized using ImageNet channel statistics, and processed with a soft Gaussian center-emphasis mask to prioritize morphology near the labelled cell while retaining local tissue context. Training augmentation included random horizontal and vertical flips, 90° rotations, and mild color jitter. Validation and test patches used the same deterministic preprocessing without stochastic augmentation.

Each backbone was trained with AdamW in two phases^46^: classification-head warm-up followed by full-model fine-tuning with cosine learning-rate scheduling and early stopping based on validation balanced accuracy. The best checkpoint for each backbone was selected by validation balanced accuracy and evaluated on the held-out test set using balanced accuracy, overall accuracy, per-class performance, and row-normalized confusion matrices. ViT-B/16 was selected for whole-slide inference.

Whole-slide motif inference. The selected ViT-B/16 classifier was applied to TCGA colorectal cancer diagnostic H&E whole-slide images using sliding-window inference. For each slide, microns-per-pixel was read from slide metadata, with objective power used as a fallback when needed. Non-overlapping windows were tiled across tissue at an effective field of view of approximately 40 µm. Each window was processed using the same deterministic validation pipeline as training patches, and the model argmax was recorded as one of four motif macro-classes. This produced a two-dimensional slide-level class map on a 40-µm grid, with each grid position assigned as DFB, Immune, VBMS, or Tumor.

Image-derived spatial biomarkers. From each predicted class map, we computed slide-level spatial biomarkers designed to quantify DFB barrier architecture and immune spatial organization. These included class fractions for DFB, Immune, VBMS, and Tumor; DFB-Tumor interface, defined as the fraction of the tumor border directly adjacent to DFB; DFB continuity, defined as the largest connected DFB component as a fraction of all DFB pixels; DFB thickness, defined from distance-transform measurements around DFB regions; and immune exclusion, defined to capture the relative spatial separation of immune pixels from the tumor in relation to the DFB barrier.

We also computed a DFB-barrier composite score as a two-component summary of DFB abundance and immune exclusion, defined as the mean of standardized DFB fraction and the negated standardized immune-exclusion term, so that higher values indicate abundant DFB with immune cells excluded from the tumor-proximal region. Slide-level biomarkers were averaged to the patient level before merging with clinical data.

#### Survival analysis

Survival analyses were performed to evaluate the prognostic associations of both transcriptomic signatures and pathology image-derived biomarkers. For gene-expression-based analyses, transcriptomic and clinical survival data from The Cancer Genome Atlas (TCGA) were accessed through the UCSC Xena platform using the UCSCXenaShiny^47^ R package. TCGA-COAD and TCGA-COADREAD cohorts were used according to the corresponding signature analysis. Overall survival time and survival status were used as the clinical endpoints.

For each predefined multi-gene signature, expression values were retrieved for each TCGA sample and merged with age and survival information by sample ID. Samples with missing or non-finite expression, age, survival time, or survival status values were excluded, and only samples with complete expression data for all genes in the corresponding signature were retained. A composite signature score was calculated for each sample as the mean expression value of the genes included in the signature. Patients were analyzed in three groups: all patients, early-onset patients defined as age < 50 years, and average-onset patients defined as age ≥ 50 years. Within each analysis group, patients were stratified into high- and low-signature groups using the median signature score as the cutoff. Kaplan-Meier survival curves were generated, and survival differences between groups were assessed using the log-rank test. Hazard ratios and 95% confidence intervals were estimated using univariate Cox proportional hazards regression. These analyses and visualizations were performed in R using the survival and survminer packages.

For pathology image-derived biomarker analyses, overall survival time and event status were harmonized from the TCGA-COAD clinical patient file and the v1.0 follow-up file by using the most recent available contact or death information for each patient. Patients were retained when both overall survival information and image-derived biomarker measurements were available. Analyses were performed in the full TCGA CRC cohort and in a pre-specified advanced-stage subset defined as AJCC stage III/IV disease.

Continuous pathology biomarkers were standardized to one standard deviation and tested for association with overall survival using Cox proportional hazards regression. Hazard ratios were reported per 1-SD increase with 95% confidence intervals. Cox models were fitted separately for each biomarker in the full cohort and in the stage III/IV subset. Within each cohort-level biomarker screen, p-values were corrected for multiple testing using the Benjamini-Hochberg false discovery rate procedure.

Kaplan-Meier survival curves were generated for selected image-derived biomarkers shown in Figure 5. Immune exclusion was stratified by median split, and the DFB-barrier composite was stratified by tertiles in the advanced-stage cohort. Group differences were assessed using the log-rank test for two-group comparisons and the multivariate log-rank test for tertile-based comparisons. To assess robustness of the advanced-stage DFB-barrier composite association, bootstrap resampling was performed with 1,000 replicates of the stage III/IV Cox model, summarizing the bootstrap hazard-ratio distribution and the probability that HR > 1.

#### Statistical analysis

Analyses were performed in Python 3.11 and R 4.3.1. Python analyses used scanpy (v1.12), squidpy^48^ (v1.8.1), AnnData (v0.12.10), NumPy (v2.2.6), pandas (v2.3.3), SciPy (v1.16.3), statsmodels (v0.14.6), scikit-learn (v1.8.0), PyTorch (v2.5.1), and PyTorch Geometric (v2.7.0). R analyses used R (v4.3.1), survival (v3.8.3), survminer (v0.5.0), ggplot2 (v3.5.2), dplyr (v1.1.4), lme4 (v1.1.37), and UCSCXenaShiny (v2.1.0). Gene ontology analysis were performed using GOrilla^49^. Benchmark presents five graph-method contrasts from this 5-peer multiplicity family.

## Figure and Table Legends

Sup Table 1. The 10-motif STORM atlas of the colorectal cancer microenvironment.

Sup Table 2. Xenium data detail information.

## Acknowledgment

We acknowledge the Whole Brain Microscopy Facility RRID:SCR_017949 for slide scanning. We acknowledge the contributions of the Tissue Management Shared Resource of the Simmons Comprehensive Cancer Center. We acknowledge the assistance of the University of Texas Southwestern Tissue Management Shared Resource, a shared resource at the Simmons Comprehensive Cancer Center, which is supported in part by the National Cancer Institute under award number P30 CA142543.

EHH was supported by NCI U01 CA214300 and NCI R01 237304. MHB was supported by the Burroughs-Wellcome Trust. This study is partially supported by the National Institutes of Health (R01GM140012 [GX], R01GM115473 GX], R01DE030656 [GX], U01CA249245 [GX]) and the Cancer Prevention and Research Institute of Texas (CPRIT RP230330 [GX]).

## Author contributions

J.Y., Y.Y., Y.J., Q.Z. contributed to the bioinformatics analyses. E.H., M.B. contributed to the generation of the CRC Xenium data. B.Y., contributed to data management. G.X., E.H., Y.X. contributed to funding acquisition. L.C.,W.S., Z.C., P.Q. contributed critical insights to the study. G.X., E.H., Y.X supervised the overall study.

## Data and code availability

All data and code will be released upon acceptance.

---

**Sup Fig 1.**
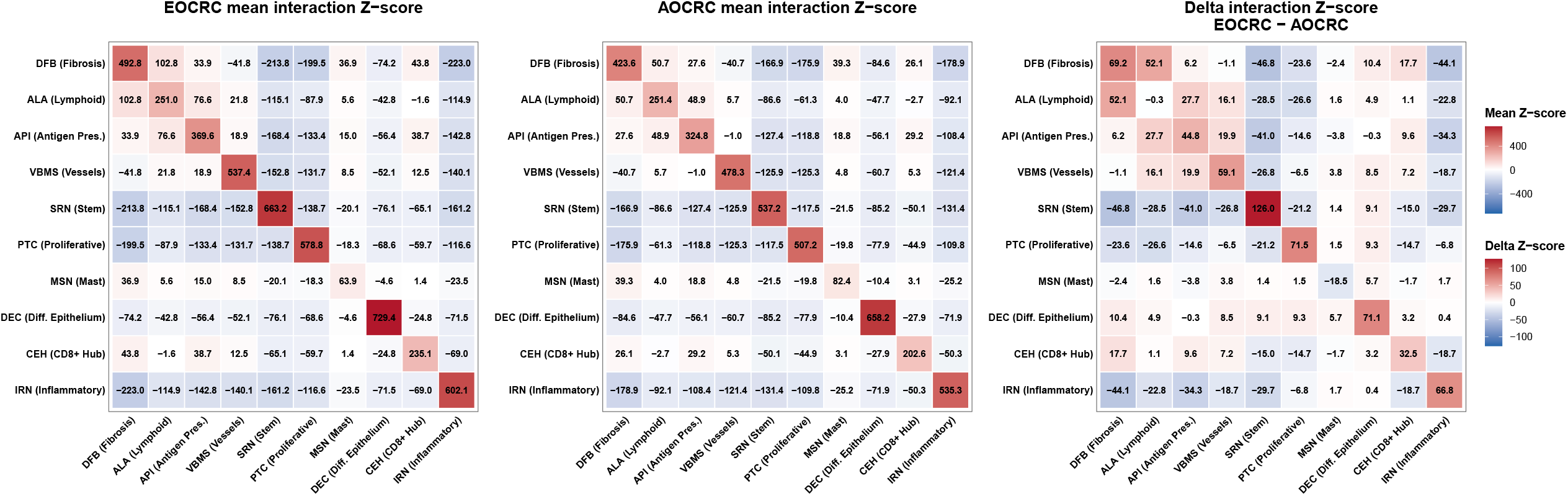
STORM model stability and robustness validation. (A) UMAP projection of per-cell STORM niche embeddings colored by motif label. (B) The same UMAP projection colored by patient, used to assess patient-driven or section-driven structure. (C) Spatial graph sanity check showing mean undirected graph degree versus cells per sample; point size and color represent the 99th-percentile edge length, and the annotation reports the total number of isolated nodes. (D) Pairwise Pearson correlation of patient-level motif-frequency profiles across the retained clustering seeds after centroid-based motif-label alignment. (E) Matched centroid cosine similarity across retained clustering seeds after Hungarian alignment to the reference seed. Points show mean cosine similarity, and vertical ranges show lower-tail to median centroid similarity summaries. (F) Leave-one-patient-out generalization, showing cosine similarity between each held-out patient’s motif-frequency vector and the mean motif-frequency vector of the training patients. (G) Pairwise embedding similarity matrix across graph-construction and receptive-field sensitivity configurations. Values show mean per-cell Pearson correlations between embedding matrices. (H) One-dimensional sensitivity profiles for k-nearest-neighbor graph size and k-hop inference depth relative to the reference configuration (k = 10, 2-hop). (I) Gene Ontology enrichment bubble plots for ALA.

**Sup Fig 2.**
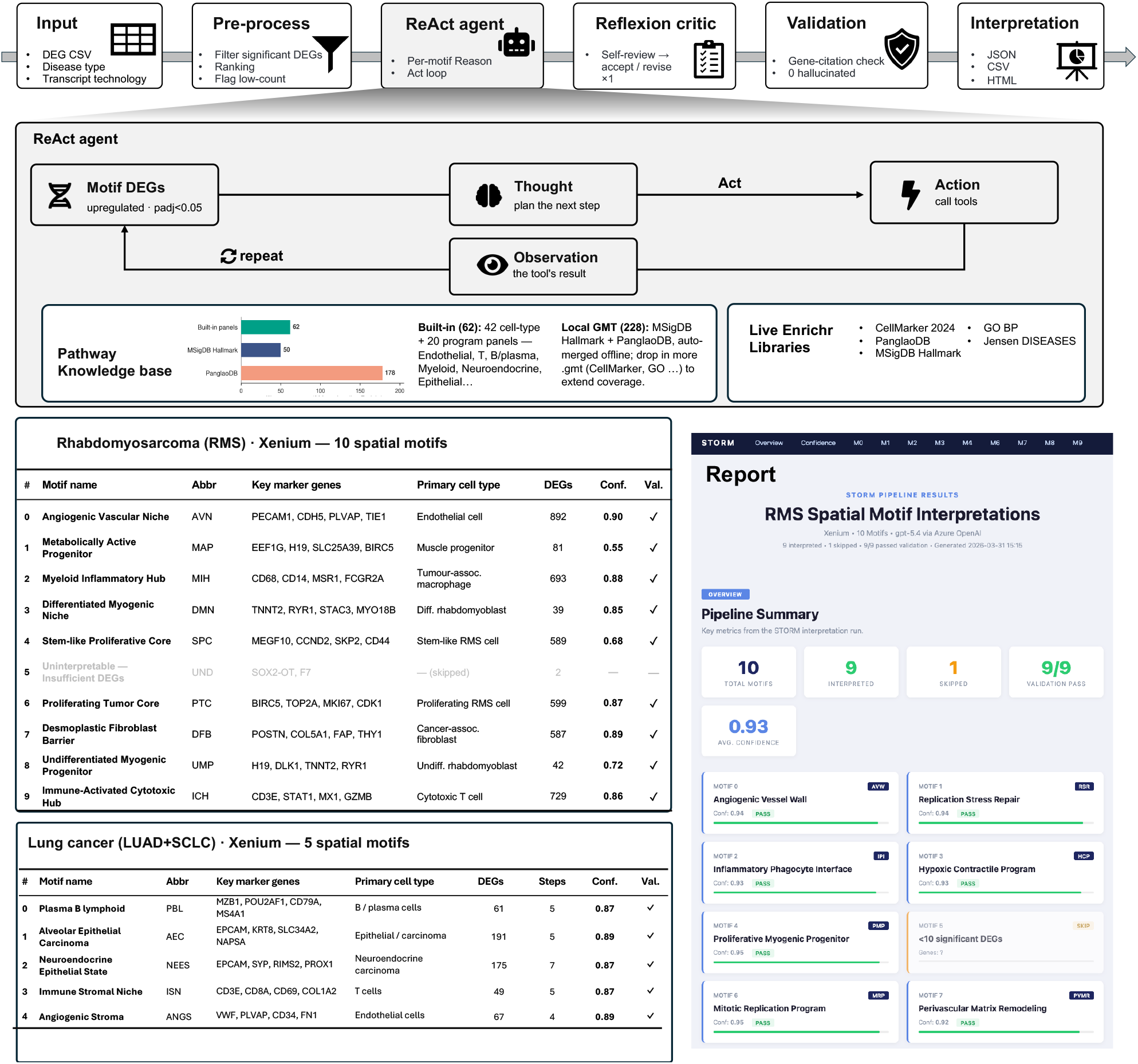
Sample-level association between DFB tumor-boundary coverage and external CD8+ T-cell access to the tumor core and penalty robustness. Each point represents one specimen, with tile-level measurements averaged within each sample (n = 16). The x-axis shows sample-level mean DFB tumor-boundary coverage, summarizing DFB boundary coverage at the specimen level. (A) Higher mean DFB coverage was associated with a higher mean barrier-crossing fraction by external CD8+ T cells (Spearman ρ = 0.503, P = 0.047). (B) Higher mean DFB coverage was also associated with an increased mean detour ratio, defined as DFB-penalized path cost divided by unweighted graph distance to the tumor core (Spearman ρ = 0.618, P = 0.011). Dashed lines indicate linear trend fits for visualization; statistical testing used sample-level Spearman correlation. (C) CD8 geodesic access robustness: penalty sensitivity (1, 3, 5, 8).

**Sup Fig 3.**
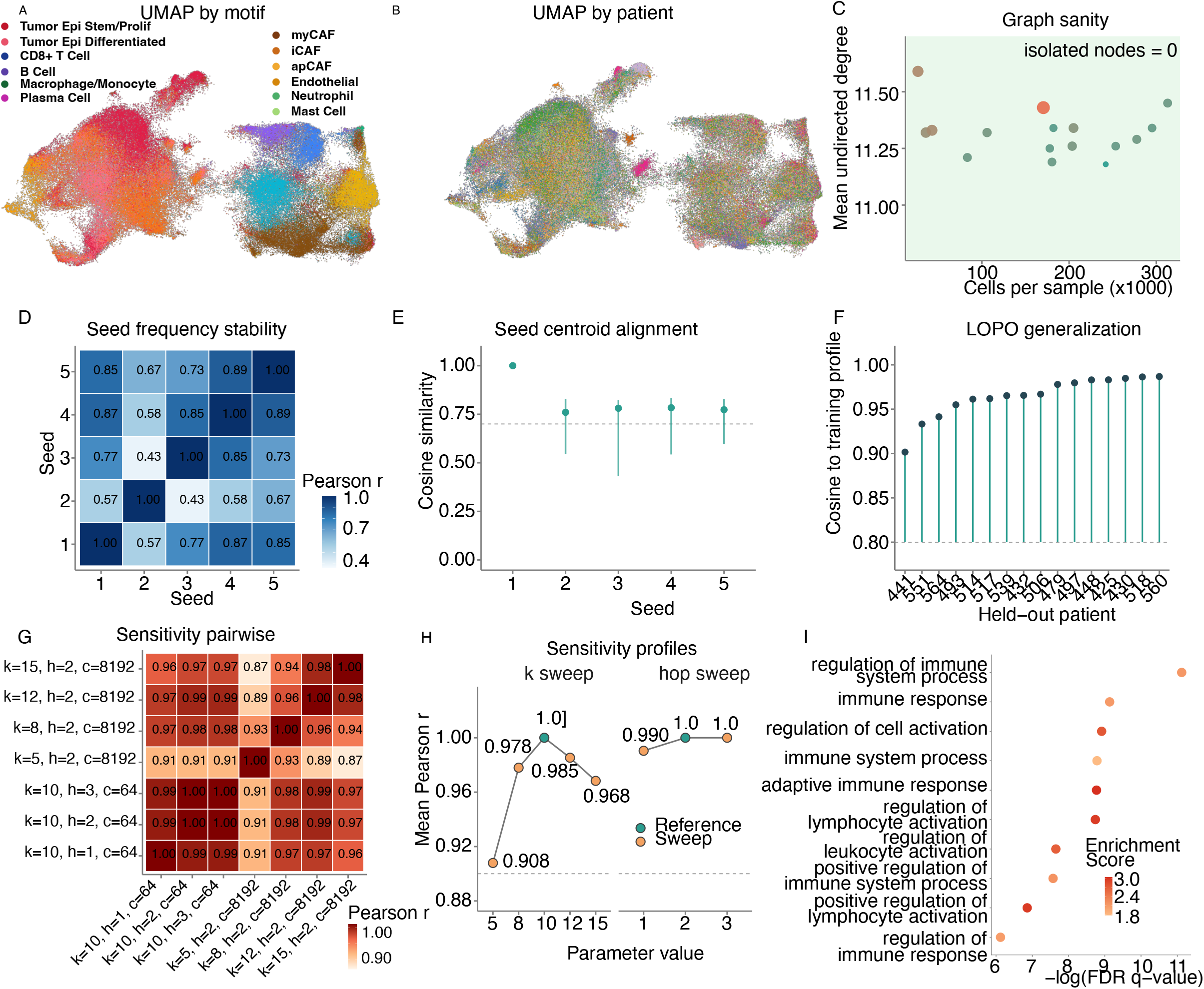
EOCRC-minus-AOCRC differential Z heatmap.

**Sup Fig 4.**
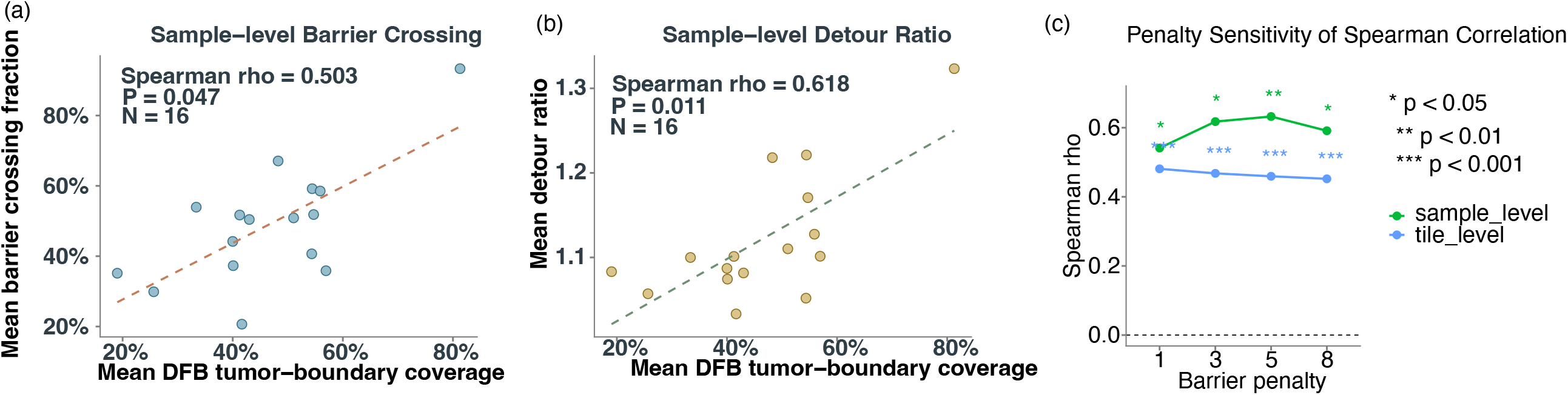
STORM-Interpret, an agentic large-language-model pipeline for spatial-motif interpretation, applied to two cancers. (A) Pipeline: STORM ingests a per-motif differential-expression (DEG) table with the disease type and transcript technology, pre-processes it by filtering significant DEGs, ranking genes, and flagging low-count motifs, and interprets each motif with a ReAct^50^ agent that iterates Thought → Action → Observation over tools spanning the motif’s statistics, an open-world pathway knowledge base (62 built-in panels + 228 offline gene sets; MSigDB Hallmark, PanglaoDB), and live Enrichr queries (CellMarker 2024, PanglaoDB, MSigDB Hallmark, GO BP, Jensen DISEASES). A Reflexion critic and a gene-citation validation step then vet the result, which is exported as JSON, CSV, and HTML. (B) Case study 1, rhabdomyosarcoma (RMS; Xenium, 10 spatial motifs): for each motif, the name, abbreviation, key marker genes, putative primary cell type, number of significant DEGs, confidence score, and gene-citation validation (✓) are listed (9 interpreted; 1 skipped for insufficient DEGs). (C) Case study 2, lung cancer (LUAD+SCLC; Xenium, 5 spatial motifs), additionally listing the number of agent tool steps per motif. (D) Auto-generated interactive HTML report for the RMS run, summarizing pipeline-level metrics (total, interpreted, and skipped motifs; validation pass rate; mean confidence) and per-motif interpretation cards. DEG, differentially expressed gene; padj, Benjamini-Hochberg-adjusted P value.

