## Supplementary material for "Spatial Topology Reveals Biologically Distinct Recurrent Motifs in Colorectal Cancer": Sup. File 1

### Quality-Control and Sensitivity Analyses for STORM Motif Discovery

#### Overview

This supplementary file describes the quality-control, stability, and sensitivity analyses used to validate the STORM spatial motif vocabulary. All analyses were performed on the per-cell niche embeddings, defined as 20-dimensional latent representations produced by the GATv2 variational graph autoencoder, and on MiniBatch k-means motif assignments derived from those embeddings. Trained model weights were held fixed for all post hoc quality-control and sensitivity analyses. The final visual summary is provided in Sup Fig 1.

#### Quality-Control and Sensitivity Analysis Methods

**UMAP and PCA Projection of Niche Embeddings.** To confirm that the discovered motifs occupied coherent regions of the learned niche-embedding space, per-cell embeddings were subsampled to a fixed number of cells per sample using a fixed random seed, keeping the point count tractable across the full Xenium cohort. The embeddings were projected to two dimensions with UMAP using umap-learn with n_neighbors = 30, min_dist = 0.3, Euclidean distance, and a fixed random seed. In parallel, the first two principal components were computed as a linear reference projection. The resulting two-dimensional embeddings were colored by motif label. The same UMAP and PCA projections were also colored by patient and sample to screen for patient-specific or section-specific structure. This batch-screening step was used to evaluate whether motif identity reflected shared tissue-niche biology rather than technical artifacts from individual patients or tissue sections.

**Spatial Graph Sanity Checks.** Each per-sample spatial neighborhood graph was constructed as a k-nearest-neighbor graph with k = 10, with edges weighted by inverse intercellular distance. Graphs were symmetrized to undirected graphs using torch_geometric.utils.to_undirected. For each sample, node degree was summarized by the mean undirected degree, maximum degree, and the number of isolated degree-zero nodes. The expected mean degree after symmetrization was between k and 2k. Edge geometry was evaluated using the Euclidean distance between connected cells from the stored spatial coordinates. Mean, median, 99th-percentile, and maximum edge lengths were summarized per sample to verify that graph edges linked physically proximal cells and were not dominated by spurious long-range connections.

**Clustering Seed Stability.** Reproducibility of motif assignments with respect to clustering initialization was evaluated by rerunning MiniBatch k-means with a batch size of 4,096 on the fixed niche embeddings at the selected number of motifs. Pairwise agreement between seed solutions was measured using adjusted Rand index and normalized mutual information. In addition, patient-level motif-frequency profiles were computed for each seed as row-normalized patient-by-motif matrices and compared across seeds after label alignment, allowing downstream patient-level motif abundance stability to be assessed directly.

**Seed Centroid Alignment.** To establish consistent motif identities across clustering runs, k-means cluster centroids from each seed were matched to those of a reference seed by maximizing total cosine similarity between centroid sets. Optimal matching was performed with the Hungarian algorithm using scipy.optimize.linear_sum_assignment on a cost matrix defined as one minus centroid cosine similarity. The resulting permutation was used to relabel motif assignments into a common reference frame before patient-level frequency profiles were compared across seeds. Mean, median, 10th-percentile, and minimum matched-centroid cosine similarities were summarized for each retained seed.

**Leave-One-Patient-Out Generalization.** Cross-patient generalizability of the motif vocabulary was evaluated by leave-one-patient-out re-clustering. For each patient, MiniBatch k-means was fit on the niche embeddings from all remaining patients using a fixed random seed. The held-out patient's cells were then assigned to the learned motif centroids by nearest-centroid prediction. The held-out patient's motif-frequency vector was compared with the mean motif-frequency distribution of the training patients by cosine similarity.

**Graph-Construction and Receptive-Field Sensitivity.** Robustness of the niche embeddings to graph-construction and inference-depth choices was assessed by reinferring embeddings from the fixed trained model under a sweep of k-nearest-neighbor values and k-hop receptive-field depths. The k-nearest-neighbor sweep used $k = 5$, 8, 10, 12, and 15, with 2-hop inference fixed. The hop sweep used 1-, 2-, and 3-hop contexts with $k = 10$ fixed. The reference configuration was $k = 10$ and 2-hop inference. For each configuration pair, the mean per-cell Pearson correlation between the two embedding matrices was computed after averaging across embedding dimensions, yielding a symmetric pairwise similarity matrix over all k and hop settings. One-dimensional stability profiles were also computed by comparing each k-sweep or hop-sweep configuration against the reference configuration. Configurations were considered stable when the mean Pearson correlation to the reference exceeded 0.95. The hop sweep was run with a small per-chunk node block so that 1-, 2-, and 3-hop contexts remained distinct during inference.

### Results

**Embedding Structure and Batch Assessment**. UMAP projection of the fixed niche embeddings showed that motif-labeled cells occupied coherent regions of the learned manifold (Supplementary Fig. 1A). Coloring the same projection by patient showed broad intermixing rather than a patient-dominant partition, supporting the interpretation that motif identity reflects shared tissue-niche biology rather than obvious patient- or section-driven batch structure (Supplementary Fig. 1B).

**Spatial Graph Quality Control.** Per-sample graph audits showed the expected topology for symmetrized $k = 10$ spatial graphs, with mean undirected degree remaining within the expected range and no isolated nodes detected across the cohort (Supplementary Fig. 1C). Edge-length summaries further supported local spatial connectivity, indicating that the neighborhood graph was not dominated by spurious long-range links.

**Seed Stability and Motif Alignment.** After centroid-based label alignment, patient-level motif-frequency profiles were positively correlated across the five retained clustering seeds (Supplementary Fig. 1D). Matched centroid similarities were also high across non-reference seeds, indicating that motif identities were preserved even when individual cluster boundaries varied modestly across initializations (Supplementary Fig. 1E).

**Cross-Patient Generalization.** Leave-one-patient-out re-clustering showed that each held-out patient's motif-frequency profile remained similar to the motif distribution learned from the remaining patients (Supplementary Fig. 1F). All 16 held-out patients exceeded the visual reference threshold used in the panel, supporting cross-patient generalizability of the motif vocabulary.

**Parameter Sensitivity and Functional Context.** Pairwise embedding comparisons across graph-construction and receptive-field settings remained highly concordant, indicating that the learned niche embeddings were not driven by a narrow hyperparameter choice (Supplementary Fig. 1G). In the one-dimensional sweeps, all k values of 8 or greater exceeded the pre-specified stability threshold, and 1-, 2-, and 3-hop inference produced highly similar embeddings relative to the reference configuration (Supplementary Fig. 1H).

Together, these analyses support the robustness of the STORM motif vocabulary: motif structure is coherent in the learned embedding space, not visibly driven by patient identity, supported by plausible spatial graph topology, stable across retained clustering seeds, generalizable across held-out patients, and robust to the principal graph-construction and neighborhood-depth hyperparameters.
