## Supplementary material for "Spatial Topology Reveals Biologically Distinct Recurrent Motifs in Colorectal Cancer": Sup File 2

Sup File 2: LLM-Assisted Spatial Motif Interpretation

### Overview

This supplementary file describes STORM-Interpret, an agentic large-language-model (LLM) pipeline for accelerating the biological interpretation of unsupervised spatial motifs discovered by STORM. The pipeline operates exclusively after motif discovery and differential expression analysis, serving as an interpretation layer that does not influence motif definition. We applied STORM-Interpret to two independent Xenium 5000-gene datasets — rhabdomyosarcoma and lung cancer — and report its annotation outputs, quality-control mechanisms, and cross-tumor observations.

### Rationale

Unsupervised spatial motif discovery produces clusters of cells defined by joint transcriptional and topological features, but assigning biological meaning to each motif traditionally requires manual inspection of marker genes, enrichment results, and spatial maps. This step is labor-intensive and scales poorly as the number of motifs and datasets grows. STORM-Interpret automates first-pass motif annotation by providing an LLM agent with structured differential-expression evidence and a curated knowledge base, while enforcing gene-citation validation to prevent hallucination. The pipeline is not intended to replace expert annotation; rather, it accelerates the transition from unsupervised clusters to interpretable biological hypotheses that can be refined or overridden by domain specialists.

### Methods

**Pipeline architecture.** STORM-Interpret is built on GPT-5.4 (Azure OpenAI, Responses API v2025-04-01-preview; temperature 0.2). The pipeline ingests a per-motif differential-expression (DEG) table together with the disease type and transcript technology, and produces structured annotations for each motif.

**Pre-processing.** The pipeline first filters the DEG table to retain only genes significantly upregulated in the motif (Benjamini-Hochberg-adjusted $P < 0.05$ and log2 fold change $> 0$), ranked by their differential score. Motifs with fewer than ten such genes are flagged and skipped, as insufficient evidence precludes reliable interpretation.

**ReAct agent loop.** Each remaining motif is interpreted by a ReAct (Reason + Act) agent that iterates through Thought-Action-Observation cycles for up to eight steps. In each step, the agent returns a single JSON object containing a thought, an action, and the action's input. The action invokes one of seven tools, and the tool's output becomes the next observation. The seven tools are:

I. Per-gene statistics: report the motif's own differential-expression statistics for queried genes.
II. Knowledge-base scoring: score the motif's genes against a curated knowledge base of cell-type and pathway panels.
III. DEG verification: check whether a named gene is a genuine DEG of the motif.
IV. Cross-motif comparison: compare the current motif with others already interpreted in the run.

Additional tools for enrichment queries, panel lookups, and summary generation.

**Knowledge base.** The knowledge base is open-world, holding 62 built-in panels and 228 offline gene sets (including MSigDB Hallmark, PanglaoDB). When built-in panels are insufficient, one tool runs a live enrichment query through Enrichr (CellMarker 2024, PanglaoDB, MSigDB Hallmark, GO Biological Process, Jensen DISEASES), falling back to the local knowledge base when offline. The knowledge base can be extended with additional files.

### Quality control

**Gene-citation validation.** The agent may cite only genes that the tools confirm are DEGs of the motif. Before the agent can finish, it must have queried the knowledge base and verified every gene it cites. Any draft that cites a non-DEG is rejected.
Reflexion critic. After the agent produces an accepted interpretation, a Reflexion critic reviews it once and triggers a single revision if needed. This step catches logical inconsistencies, unsupported claims, or missed alternative interpretations.
Final validation. A final validation step re-checks all gene citations against the input DEGs. Only interpretations that pass this step are retained.

**Output format.** For each motif, STORM-Interpret returns:

- Motif name and abbreviation
- Key marker genes (validated DEGs only)
- One or more putative cell types
- Confidence score
- Short gene-cited description (<=150 words)
- Brief limitations note (<=50 words)

All outputs are saved as JSON, a summary CSV, an interactive HTML report, and the full reasoning trace.

### Application datasets

We applied STORM-Interpret to two independent Xenium 5000-gene datasets on which STORM had been run separately to discover spatial motifs without cell-type labels or manual annotations:
(i) Rhabdomyosarcoma (RMS): 10 spatial motifs. (ii) Lung cancer (LUAD + SCLC): 5 spatial motifs.
In each dataset, STORM first learned topology-aware cell embeddings and defined spatial motifs. Differential expression analysis was then performed for each motif, and the resulting DEG tables were provided to STORM-Interpret for biological interpretation.

### Results

**Rhabdomyosarcoma.** Nine of the ten RMS motifs met the evidence threshold (>=10 significant DEGs) and were interpreted by the agent; one motif was skipped due to insufficient DEGs. The agent generated structured annotations including proposed motif names, likely biological functions, supporting marker evidence, alternative interpretations, and confidence scores. The nine interpreted motifs were organized into tumor, stromal, immune, vascular, and tissue-remodeling programs ( Sup Fig. 4B).
Notably, one stromal motif was annotated as a fibroblast- and extracellular-matrix-rich desmoplastic barrier-like structure, characterized by collagen and matrix-remodeling programs. This motif resembles the Desmoplastic Fibrotic Barrier (DFB) architecture identified in our CRC cohort.

**Lung cancer.** All five lung cancer motifs met the evidence threshold and were interpreted. The agent organized the motifs into interpretable tumor, stromal, immune, and vascular programs (Sup Fig. 4C). As in the RMS dataset, a fibroblast- and ECM-rich stromal motif was identified and annotated as a desmoplastic barrier-like structure with collagen and matrix-remodeling signatures.

**Cross-tumor DFB-like motif observation.** In both independent datasets, STORM-Interpret identified a stromal motif with transcriptional features consistent with the DFB architecture observed in our colorectal cancer cohort — specifically, enrichment for fibrillar collagens, matricellular proteins, and matrix-remodeling enzymes. Although this observation is exploratory and based on transcriptional signature alone, it suggests that DFB-like spatial architectures may recur across tumor types beyond colorectal cancer.

### Limitations

We emphasize the following caveats:

1. The DFB-like annotation is based on transcriptional signature only. Spatial topology analyses — including neighborhood-enrichment Z matrices, interface/non-interface DFB partitioning, and graph-geodesic barrier modeling — were not performed on the RMS or lung cancer datasets. Whether these DFB-like motifs exhibit the same layered barrier-exclusion topology observed in CRC remains to be tested.
2. LLM-generated annotations are interpretive hypotheses, not ground truth. They are subject to the limitations of the underlying LLM, the knowledge base, and the DEG evidence, and should be validated by domain experts.
3. The pipeline depends on a specific LLM version (GPT-5.4); outputs may change with model updates, though the gene-citation validation step constrains the space of admissible annotations.
